# Thermal imaging can reveal variation in stay-green functionality of wheat canopies under temperate conditions

**DOI:** 10.1101/2023.11.08.566006

**Authors:** Jonas Anderegg, Norbert Kirchgessner, Helge Aasen, Olivia Zumsteg, Beat Keller, Radek Zenkl, Achim Walter, Andreas Hund

**Affiliations:** Plant Pathology Group, Institute of Integrative Biology, ETH Zurich, Zurich, Switzerland; Crop Science Group, Institute of Agricultural Sciences, ETH Zurich, Zurich, Switzerland; Earth Observation of Agroecosystems Team, Research Division Agroecology and Environment, Agroscope, Zurich, Switzerland

**Keywords:** high throughput field phenotyping, physiological breeding, deep learning, semantic segmentation, remote sensing

## Abstract

Canopy temperature (CT) is often interpreted as representing leaf activity traits such as photosynthetic rates, gas exchange rates, or stomatal conductance. Accordingly, CT measurements may provide a basis for high throughput assessments of the productivity of wheat canopies during early grain filling, which would allow distinguishing functional from dysfunctional stay-green. However, whereas the usefulness of CT as a fast surrogate measure of sustained vigor under soil drying is well established, its potential to quantify leaf activity traits under high-yielding conditions is less clear. To better understand sensitivity limits of CT measurements under high yielding conditions, we generated within-genotype variability in stay-green functionality by means of differential short-term pre-anthesis canopy shading that modified the sink:source balance. We quantified the effects of these modifications on stay-green properties through a combination of gold standard physiological measurements of leaf activity and newly developed methods for organ-level senescence monitoring based on timeseries of high-resolution imagery and deep-learning-based semantic image segmentation. In parallel, we monitored CT by means of a pole-mounted thermal camera that delivered continuous, ultra-high temporal resolution CT data. Our results show that differences in leaf activity stemming from differences in stay-green functionality translate into measurable differences in CT in the absence of major confounding factors. Differences amounted to approximately 0.8°C and 1.5°C for a very high-yielding source-limited genotype, and a medium-yielding sink-limited genotype, respectively. The gradual nature of the effects of shading on CT during the stay-green phase underscore the importance of a high measurement frequency and a time-integrated analysis of CT, whilst modest effect sizes confirm the importance of restricting screenings to a limited range of morphological and phenological diversity.

## Introduction

The onset of monocarpic senescence is a critical phenological event in annual crops, marking a basic transition of canopies from carbon assimilation to Nitrogen remobilization (Thomas and Ougham, 2014). Senescence-related remobilization processes are pivotal for yield and quality formation in wheat (Kichey et al., 2007), but an adequate post-anthesis green canopy duration preceding its onset is similarly important to avoid time- and resource-related constraints to grain filling. Indeed, maximizing carbon assimilation by a prolonged maintenance of green leaf area after anthesis (the “stay-green” trait) represents an important breeding aim in several crops (reviewed by Gregersen et al., 2013).

Single genes with major effects on the timing and dynamics of senescence have been identified in wheat (Uauy et al., 2006), but senescence is generally considered to be under complex genetic and environmental control. Numerous studies highlighted the importance of the balance between Nitrogen uptake from the soil and Nitrogen demand by developing grains as a determinant of its timing and dynamics (Kichey et al., 2007; Rajcan and Tollenaar, 1999; Triboi and Triboi-Blondel, 2002; van Oosterom et al., 2010). From this perspective, a timely and rapid senescence indicates a high sink demand for assimilates and Nitrogen (Xie et al., 2016; Yang and Zhang, 2006), whereas an unfavorably delayed and slow senescence indicates sink-source imbalances, such as resulting from a low yield potential or overfertilization (Jiang et al., 2004; Naruoka et al., 2012; Yang and Zhang, 2006). When selecting for increased green canopy duration, it would therefore be imperative to distinguish between functional stay-green associated with enhanced photosynthetic activity required to meet a high sink demand, and dysfunctional stay-green resulting from slowed Nitrogen remobilization indicating low sink demand and, consequently, a low yield potential (Gregersen et al., 2008). This may be particularly important under high yielding conditions where strong external environmental triggers of senescence such as heat or drought events are absent in average years.

Key aspects of senescence and its dynamics, such as rates of chlorophyll degradation, are readily observable by the eye, both at the canopy level as well as at the level of visible parts of plant organs (e.g., leaves, peduncles, or ears). Numerous studies have used proximal sensing techniques based on average canopy light reflectance to track the dynamics of senescence (e.g., Anderegg et al., 2020; reviewed by Chapman et al., 2021). High-resolution imaging combined with deep-learning-based semantic image segmentation additionally enabled tracking of senescence and overall healthiness of wheat stands at the organ-level (Anderegg et al., 2023). In comparison to visual assessments, these approaches offer the key advantages of objectivity and scalability. Unfortunately, however, much like visual assessments, they do not enable a distinction between functional and dysfunctional stay-green, nor a precise assessment of remobilization, grain filling rates or grain filling duration, since an increased green canopy duration as observed visually is not per se indicative of increased grain filling rates or duration. Therefore, measurements of canopy greenness must be complemented with measurements of “leaf activity” traits (Fischer et al., 1998), such as photosynthetic activity, gas exchange rates, or stomatal conductance.

Numerous retrospective studies on historical series of genetic lines have found remarkably strong correlations between increases in stomatal conductance and yield gains over time (i.e., with year of release; e.g., Fischer et al., 1998; reviewed by Roche, 2015). Increased sink-to-source ratios are hypothesized to contribute significantly to this relationship (Roche, 2015). With all else equal, higher transpiration rates (a greater stomatal conductance) decrease canopy temperature (CT) *via* evaporative cooling; therefore, CT measurements have been used as a fast surrogate measure for stomatal conductance (e.g., Rebetzke et al., 2013). Additionally, since photosynthetic gas exchange and stomatal conductance are intertwined, CT may provide an indirect measurement of photosynthetic rate (Amani et al., 1996; Fischer et al., 1998; Jones and Vaughan, 2011). Recently, CT measurements have been acquired using airborne imaging thermography, enabling the measurement of large experiments in short time, which greatly increased repeatability of measurements as compared to plot-by-plot measurements using hand-held thermometers (e.g., Deery et al., 2019, 2016; Perich et al., 2020).

While these studies represent significant methodological advances and provide convincing evidence for the potential usefulness of airborne CT measurements in breeding, some unexpected patterns also became apparent, including in our own data, which was collected using repeated drone-based thermal imaging of large germplasm collections throughout the grain filling phase (Anderegg et al., 2021; Perich et al., 2020). Most notably, a high heritability of plot-based CT was observed even at maturity, when no transpiring leaf tissue was left. Additionally, CT at maturity was moderately to highly correlated with CT shortly after flowering, as well as with CT measured throughout grain filling. These correlations were comparable in magnitude to correlations observed between CT values at earlier measurement dates during the stay-green phase (see for example Figures 8 and 9 in Perich et al., 2020) and this cannot be well explained when interpreting CT primarily as a measure of leaf activity traits. Finally, but perhaps less surprisingly, moderate to strong correlations were observed between CT and structural, morphological, and phenological characteristics of genotypes (Anderegg et al., 2021), which is in line with results from numerous other studies (see reviews by Deery and Jones, 2021 and Prashar and Jones, 2014). Taken together, these observations prompted some skepticism on our side regarding the existence of a direct and strong enough link between leaf activity traits and remotely sensed CT under the conditions of the study site (i.e., high-yielding zones of temperate Europe). This is mentioned here not to question the usefulness of CT as a valuable tool in breeding, for which ample evidence has been presented by others (e.g., Li et al., 2019; Lopes and Reynolds, 2010; Rebetzke et al., 2013; Thapa et al., 2018). However, we want to highlight the need for a better understanding of the extent to which CT measured at different growth stages and under different growing conditions can be interpreted as representing directly leaf activity traits. We anticipate that such an improved understanding will help quantify the value of CT measurements as a complement to precision assessments of phenology and measurement of canopy biophysical characteristics in the characterization of the status of stay-green canopies.

Therefore, the objective of this study was to isolate and quantify the direct effect of differences in leaf activity traits on remotely sensed CT. In other words, we aimed to establish whether differences in photosynthetic rates and stomatal conductance are detectable in canopies differing only with respect to these traits, but with all else as similar as possible and growing side-by-side. To this end, we aimed to introduce variation in terms of the functionality of stay-green in otherwise identical canopies. We modified sink-source relationships by applying short-term canopy shading during rapid spike growth with the aim of reducing potential yield, whereas directly neighboring control plots of the same genotype were left untreated. Plot CT was continuously monitored throughout grain filling by means of a pole-mounted thermal camera, and precision phenology assessments were made using frequent RGB imaging combined with segmentation of different organs based on deep learning models. Our results clearly indicate that differences between functional and dysfunctional stay-green translate into measurable differences in CT, and that they do so in the absence of co-variation in often correlated confounding traits.

## Materials and Methods

### Plant materials, experimental design, and environmental data

A field experiment with two registered winter wheat cultivars (‘Piznair’, AGROSCOPE/DSP, Switzerland; and ‘Campesino’, SECOBRA Saatzucht GmbH, Germany) was carried out at the ETH Research Station for Plant Sciences Lindau-Eschikon, Switzerland (47.449°N, 8.682°E, 520 m above sea level; soil type: eutric cambisol) in the wheat growing season of 2022–2023. The two cultivars were selected for the experiment based on (i) their similarity in terms of canopy characteristics, particularly leaf and ear orientation and final height; (ii) their similar phenology: both cultivars are classified as mid-late in terms of ear emergence and maturation; and (iii) their strongly contrasting yield potential: ‘Campesino’ has a very high yield potential, whereas ‘Piznair’ has an intermediate yield potential but a high protein content (Strebel et al., 2022).

Each cultivar was grown in ten plots for a total of 20 experimental plots. Wheat was sown with a drill sowing machine in nine rows per plot with a row length of 1.7 m and a row spacing of 0.125 m at a density of 400 plants m^-2^ on 18 October 2022. Plots were arranged in pairs sown with the same cultivar (Figure 1C). To avoid inhomogeneous neighboring effects on experimental plots, each pair of plots was bordered both in sowing direction as well as perpendicular to it by buffer plots sown with a late-maturing check with a very similar canopy height (‘Montalbano’, AGROSCOPE/DSP, Switzerland). Pairs of plots were arranged in a cultivar-alternating manner in two buffer-separated ranges (Figure 1A, 1C). At booting (growth stage [GS] 43; reached on 13 May 2023) one plot in each pair was shaded by suspending a polyethylene shading net that decreased light intensity by 73% (Agroflor, Wolfurt, Austria) approximately 25 cm above the top of the canopy (Figure 1B), whereas the adjacent plot was used as an unshaded control. The spatial arrangement of control and shaded plots was alternated across the pairs. Shading was applied after full flag leaf emergence in order to avoid undesired side-effects on canopy characteristics such as total above ground biomass, leaf area index, canopy cover, or leaf sizes. The shading nets were removed again at the late heading stage (approximately GS 59, reached on 29 May 2023). Crop husbandry was performed according to local agricultural practice. Temperature, rainfall, and wind speed data were retrieved from an on-site weather station.

**Figure 1.**
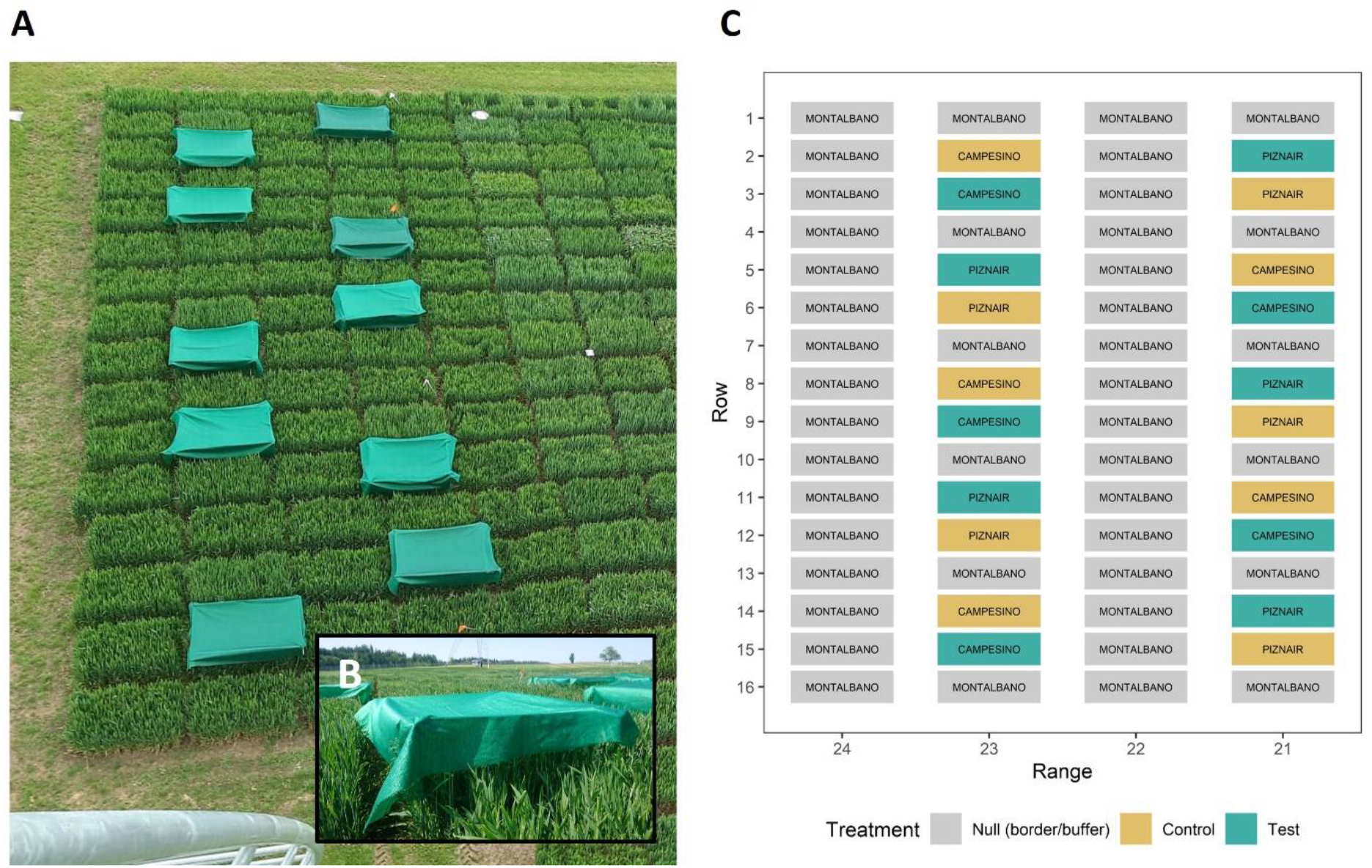
Design of the field experiment. **(A)** Image of the field experiment taken from the position where the thermal camera was mounted. The image was taken right after the shading tents were mounted on 13 May 2023 (GS 43); **(B)** close-up of a shading tent; **(C)** schematic of the experimental design. The scale of the axes was inverted to directly represent the view as seen in (A). Control plots were left untreated, whereas Test plots were shaded during rapid spike growth. Text labels within color boxes represent cultivar names. The experiment was bordered to the right side by an additional row of border plots, identical to range 24. The experiment was sown row-wise on 18 October 2022.

### Measurement of gas exchange and photosynthetic parameters

Gas exchange measurements were made plot-by-plot on eight dates between 8 June and 29 June 2023, using the portable photosynthesis system LI-6400XT (LI-COR, Inc., Lincoln, NE, USA). Measurements were made on fully sun-exposed, intact flag leaves with no visible disorders, at a position about halfway along its length. Measurements were made under clear sky conditions between 10 a.m. and 3 p.m. This ensured that leaves had been exposed to constant light prior to the measurement. One measurement lasted 3 min, with one value logged every 5 s. Air temperature in the chamber was regularly controlled and adjusted as needed to match the ambient temperature in the field. The air flow rate during the measurements was 300 µmol air s^−1^. To ensure stable CO_2_ concentration in the incoming airflow, a canister with an open cap was used as a buffer volume, as recommended by the manufacturer. Infrared gas analyzers were matched once before a measurement series. The light source intensity was set to 1500 µmol m^-2^ s^−1^. All 20 plots of the experiment were routinely measured within 1.5 h. One or two measurement runs were made on each measurement date. For each measured sample, the first 6 logged values were removed. After that, one iteration of outlier removal was performed on logged values, defining an outlier as a value deviating from the median by more than 1.5 times the interquartile range. The remaining values were averaged to obtain one value for each measured leaf. The experiment was measured twice on 8 June and 12 June 2023, and once on all other measurement dates. Whenever more than one measurement was performed, values across measurement runs were averaged on a plot basis.

Active fluorescence measurements were obtained on twelve dates between 1 June and 29 June 2023, using a MultispeQ device (PhotosynQ Inc, MI, USA). The ‘Photosynthesis RIDES 2.0_short’ protocol (photosynq.org) was used to measure the operating efficiency of photosystem II (F_q_’/F_m_’), photosynthetic photon flux rate (PPFR), and relative chlorophyll (SPAD) (Keller et al., 2023; Kuhlgert et al., 2016). Measurements were performed on leaves that were selected following the same principles as for gas exchange measurements. Between two and six measurement runs were performed on each date. Whenever more than one measurement was performed, values across measurement runs were averaged on a plot basis.

### Crop phenology, morphology, and agronomic traits

The dynamics of senescence were monitored with nearly daily resolution at the level of individual organs by means of repeated imaging from a nadir as well as an off-nadir perspective (viewing angle of approximately 45°), using a measurement setup described in detail earlier (Anderegg et al., 2023; Grieder et al., 2015). The resulting image time series were first segmented into a vegetation and a soil fraction. Subsequently, ears were segmented in images captured from a nadir perspective, whereas ears and stems were segmented in off-nadir images. Nadir images were segmented using deep convolutional neural networks available from previous work, without modification (Anderegg et al., 2023). The stem and ear segmentation model for off-nadir images was developed in the context of this study, following the same procedure as described for nadir images earlier (Anderegg et al., 2023). Briefly, 100 patches sized 600 × 600 pixels from carefully selected diverse images representing the entire grain filling phase, contrasting light conditions, and different genotypes, were manually annotated at pixel level, using polygons. This data set was complemented with 165 patches from images captured with a different setup and containing only stay-green canopies. Compared to the target domain, these images had a much higher resolution but a much shallower depth of field, resulting in a blurry background. These images were resized to match the target domain in terms of physical resolution. 24 randomly selected patches from the target domain were designated as the validation data set. Model hyper-parameters were optimized within a limited search space, in a stepwise procedure. First, the segmentation framework together with the input resolution were optimized, with resnet34 (He et al., 2016) as encoder. Next, the resnet encoder depth was optimized together with data augmentation (image resolution, rotation, horizontal and vertical flipping, down-scaling and up-scaling, color jittering, blurring), and the training strategy. Finally, parameters of the network training process were optimized. The resulting optimized segmentation model reached an overall validation F1-Score of 0.90 (see Supplementary Figure S1 for an illustration). The resulting segmentation masks enabled an estimation of the fraction of images representing different components of vegetation. The original images were further segmented pixel-wise based on color properties of pixels into a green, chlorotic, and necrotic fraction, using a previously trained classifier (Anderegg et al., 2023). The masks were combined through logical operations to obtain the fractions of green, chlorotic, and necrotic tissues for each vegetation component. For details, refer to (Anderegg et al., 2023). The annotated data sets representing the target domain will be made freely available via the Repository for Publications and Research data of ETH Zürich (https://www.research-collection.ethz.ch/).

Peduncle length was measured shortly before harvest as the distance between the uppermost node on the stem and the spike collar for 12 randomly selected culms per plot with a precision of 1 cm, using a ruler. Peduncle senescence was assessed by visually classifying them as either senescent or green. Three batches of 10 tillers were examined per plot, and the total fraction of senescent peduncles was recorded. All assessments were made as recommended by Chapman et al. (2021) and Pask et al. (2012). As peduncle senescence was used as a reference for the image-derived stem senescence score, plots in a separate experiment were additionally scored to broaden the validation basis, and off-nadir imagery was also collected in the context of that experiment.

Spikes were sampled for volume measurements on 10 June, 4 July, and 11 July 2023. On each date, two spikes were sampled for each plot. Sampled spikes were stored under dry conditions and were later scanned using a 3D light scanner (Shining 3D Einscan-SE V2, SHINING3D, Hangzou, China). Parts that were used to stabilize the spike in the scanner were removed from the resulting point cloud using a custom MatLab script (MatLab r2022b, Natick, MA, USA). The spike volume was extracted using pymeshlab (https://github.com/cnr-isti-vclab/PyMeshLab). Some strongly bent spikes had to be scanned in a special mounting. The points of these mountings were manually removed with 3dbuilder (https://www.microsoft.com/en-us/3d-print/3d-builder-users-guide?rtc=1) and the spike volume was determined using Meshlab 1.3.2_64 bit (https://www.meshlab.net/).

Grain yield and total above-ground biomass were assessed by manually harvesting the sowing rows 6, 7, and 8 (out of 9). After the ears had been removed for determination of grain yield, the remaining biomass was cut approximately 1 cm above ground and dried to constant weight. Grain protein concentration (GPC) was determined using near-infrared transmission spectroscopy (IM-9500, Perten, Hägersten, Sweden). Thousand kernel weight (TKW) was measured using a MARViN ProLine II (MARViTECH GmbH, Wittenburg, Germany).

### Thermal imaging and extraction of plot canopy temperature

A thermal camera (FLIR A655sc uncooled microbolometer camera with W/45° lens, 640 × 480 pixel; FLIR Systems AB, Sweden) was mounted on a pole of the ETH field phenotyping platform FIP (Kirchgessner et al., 2017) located right next to the experimental field (cf. Figure 1A). The camera was mounted in a weatherproof housing (Tecnovideo, Villaverla, Italy) at a height of 23.5 m above ground and connected to a 12V power supply. It was controlled by a common PC at the bottom of the pole connected by LAN. Image acquisition was controlled through a Matlab script (r2022b, The Mathworks, Natick, USA), which wrote data directly to our NAS. Viewing angles for experimental plots were between 34° and 46.5° to nadir view. The used lens provides a field of view of 45° and 34°. An image was recorded every 20 s, resulting in approximately 140.000 images that covered the entire grain filling phase from flowering to shortly before harvest. From each image, median plot temperatures were extracted by generating a geojson file that contained the corner coordinates of polygons representing each experimental plot using the ‘ogr’ module of the python library ‘osgeo’. The shapes were generated by specifying the number of rows and ranges of the experiment as well as the size of a plot, with plot length set to 1.1 m, and plot width set to 0.8 m, thus leaving a buffer zone of approximately 0.3 m in sowing direction and 0.2 m perpendicular to it. Plot shape corner coordinates were then transformed by calculating their dot product with the homography matrix, which was determined by matching all four corners of the experiment to the pixel coordinates of the respective position in one example thermal image. These steps were accomplished using code associated with Treier et al. (2023). Before exporting summary statistics per plot, outliers were removed as pixel values that deviated from the median value of all pixels attributable to that plot by more than 1.5 times the interquartile range. This was deemed necessary to reduce the effect of obstructions in images, such as for example a person performing measurements in the experiment.

### Statistical Analysis

Canopy temperature time series were smoothed by fitting a smoothing spline using the function *smooth.spline()* of the R-package ‘stats’ (R Core Team, 2018), separately for each experimental plot. The number of spline knots was set to one tenth of the number of observations, and a prediction every 2 min from the resulting fits was retained for further analyses. To summarize the resulting smoothed time series of CT measurements, we extracted the area under the curves (AUC_CT_) for the duration between 10 a.m. and 4 p.m. at each measurement date.

The experimental design resulted in a spatially perfectly homogeneous distribution of control and test plots for each of the two evaluated genotypes, and there were ten direct neighbor pairs of control and test plots (five for each cultivar), that were themselves surrounded by invariable buffer plots. We therefore considered neighboring plots as representing paired samples. For canopy temperature measurements, this accounted for variation over time attributable to short-term fluctuations in environmental conditions, as well as for spatial effects related to field heterogeneity and measurement geometry. For plot-by-plot measurements, it additionally allowed for a correction of temporal effects since test and control plots were allocated to the members of a pair in a spatially alternating fashion (Figure 1). Plot-based values were therefore compared across the treatments by means of a paired samples t-test, carried out separately for each date.

The effects of genotype, treatment, and genotype-by-treatment interactions on phenological, morphological and agronomic trait values measured at the plot level were determined through a corresponding two-way analysis of variance, conducted independently for each trait, using the R-function *aov()*. Pairwise comparisons were made by means of a Tukey posthoc test, using the function *TukeyHSD()*. Where repeat measurements were made for each plot, a linear mixed effects model was fitted, with the experimental plot additionally considered as a random effect. These models were fitted using the function *lme()* of the R-package ‘nlme’ (Pinheiro et al., 2021) and all possible pair-wise contrasts were obtained from the function *emmeans()* of the R-package ‘emmeans’ (Lenth et al., 2020).

## Results

### Pre-anthesis canopy shading effectively modified the sink-source relationship with minor effects on other canopy characteristics

Canopy shading was applied during rapid spike growth and after full flag leaf emergence with the aim of generating variation in sink strength within each tested genotype. This, in turn, was expected to create variation in leaf activity traits in response to the modified sink-source balance, in canopies with otherwise very similar characteristics.

The shading treatment drastically reduced grain yield in both genotypes, with a larger relative reduction observed for the low-yielding cultivar ‘Piznair’ (Δ = -38.4%, p<0.001; Figure 2A), than for the high-yielding cultivar ‘Campesino’ (Δ = -28.4%, p<0.001). The reduction in yield potential for each genotype was clearly apparent already in the first measurement of ear volume on 10 June 2023 (i.e., 12 d after full heading [GS 59]) in terms of a reduced spike volume, which was also more prominent in the low-yielding cultivar (Δ = -36.5%, p<0.001; Figure 2D) than in the high yielding cultivar (Δ = -30.1%, p<0.01). The difference in spike volume was mostly attributable to an increased number of rudimentary basal and apical spikelets that did not develop grains (Figure 2G, 2I, 2J). This had notable canopy-level effects, with decreased ear coverage both in nadir and in oblique images for shaded than for control canopies (Figure 2G, 2E, Supplementary Figure S2). In contrast, ear volumes were not different between genotypes within each of the treatments despite the large differences in end-of-season grain yield (p > 0.8 for both treatments; data not shown).

**Figure 2.**
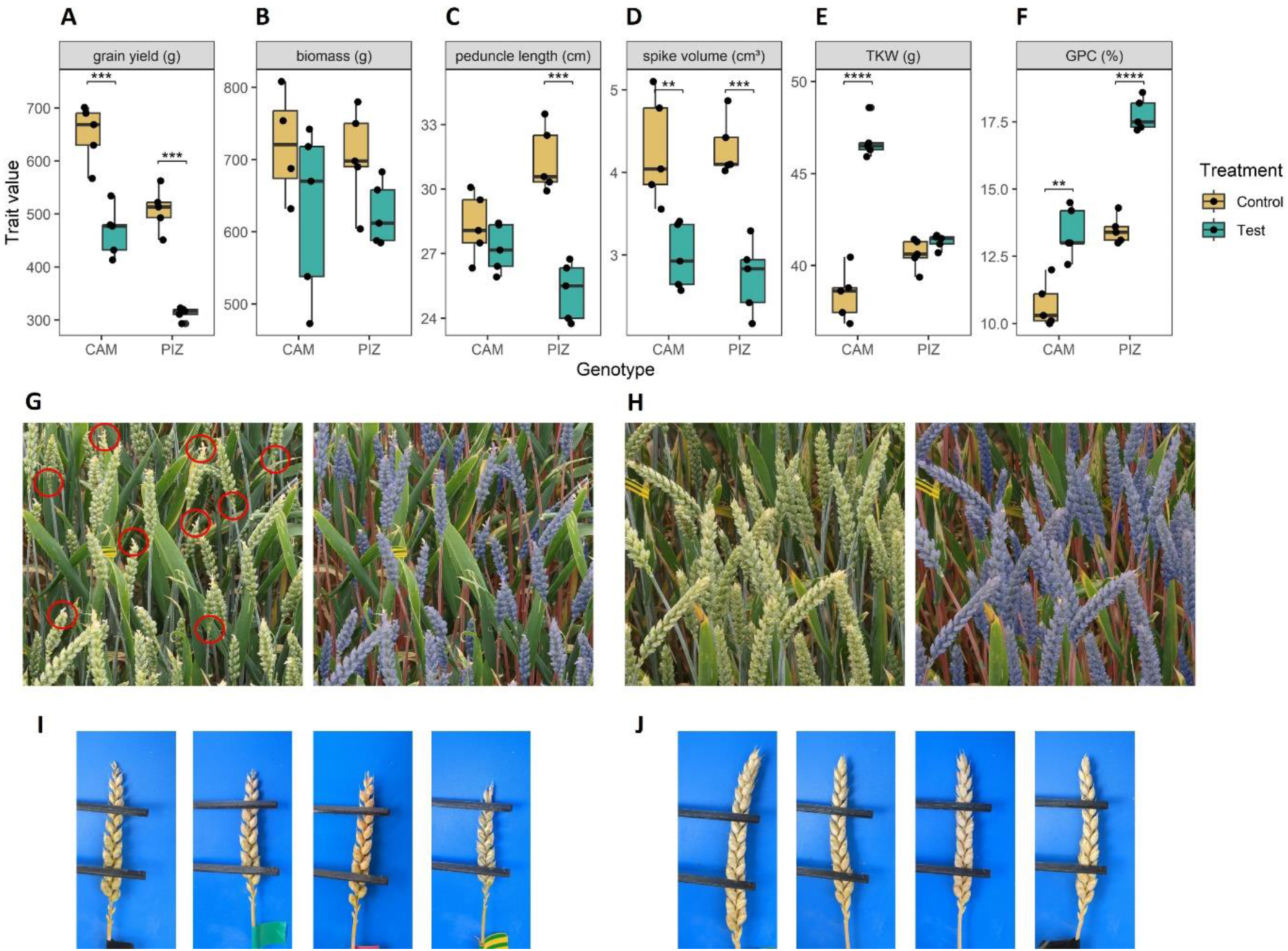
Effects of canopy shading on agronomic traits and canopy characteristics. Effects of shading on **(A)** grain yield, **(B)** above ground vegetative dry biomass (total above ground biomass after threshing), **(C)** peduncle length, **(D)** spike volume, **(E)** thousand kernel weight, **(F)** grain protein concentration. Where multiple measurements were made per plot, the mean value across repeat measurements is plotted. **(G)** Example of an image of a shaded plot with the corresponding segmentation model output overlayed (right side) for the cultivar ‘Piznair’; red circles highlight some obvious instances of rudimentary basal and completely aborted apical spikelets; **(H)** image of the directly adjacent unshaded plot. See supplementary Figure S1 for quantitative results on organ contribution to images; **(I)** Close-up images of spikes from a shaded plot of ‘Piznair’; **(J)** close-up images of spikes from the adjacent control plot of ‘Piznair’.

Thousand kernel weight was much increased under the shading treatment in ‘Campesino’ (Δ = +21.9%, p<0.001; Figure 2), indicating that grain yield in this genotype was source-limited under the control treatment; no increase in thousand kernel weight was observed for ‘Piznair’, indicating sink-limitation under the control treatment for this genotype. Grain protein concentration was increased in response to shading in both genotypes (Figure 2F). In ‘Piznair’, grain protein concentration reached a very high value of 17.8% under shading.

Despite the late application of the shading treatment, a trend towards a reduced above ground vegetative biomass was also observed, though these effects were not statistically significant (Δ = -12.8%, p=0.36 and Δ = -11.2%, p=0.44 for ‘Campesino’ and ‘Piznair’, respectively; Figure 2B). The observed trend towards a decreased vegetative biomass in shaded canopies can be partly explained by the effect of shading on peduncle length (Δ = -3.89%, p=0.64 and Δ = -19.3%, p<0.001 for ‘Campesino’ and ‘Piznair’, respectively; Figure 2C). In ‘Piznair’, the reduction in peduncle length was significantly correlated with the reduction in vegetative biomass (Pearson r = 0.66, p<0.001; not shown). In contrast, canopy cover was not affected by shading: it was nearly 100% in oblique-angle images and approximately 0.8 in nadir images, irrespective of the treatment (Supplementary Figure S2). We observed no differences in canopy characteristics besides the mentioned differences in peduncle length and ear fraction. In particular, leaf sizes and orientation appeared to be unaffected by the treatment. No differences were expected, given the late application of the treatment.

### Measurements of phenology, photosynthetic rates, and gas exchange indicated extensive dysfunctional stay-green in response to pre-anthesis canopy shading

Shading tended to delay senescence in both genotypes. In the absence of shading, the onset of senescence occurred at similar timepoints for both genotypes but progressed much faster in ‘Piznair’ than in ‘Campesino’ (Figure 3, left-most column). In both genotypes, the delay in senescence was most pronounced in leaves, but less in ears and stems (Figure 4A, 4B, 4C). As observed for agronomic traits, the delay in visible senescence in response to shading was greater for ‘Piznair’ than for ‘Campesino’ (Figure 3; Figure 4A, 4B, 4C). On average, the mid-point of foliar senescence was delayed by 2.4 d (p = 0.09) and 4.4 d (p < 0.001) in ‘Campesino’ and ‘Piznair, respectively (Figure 4C). In shaded plots of ‘Piznair’, senescence progressed much slower than in unshaded control plots, especially in leaves (Figure 3D-F). Strikingly, in shaded plots senescence progressed slower in ‘Piznair’ than ‘Campesino’, which is the opposite of what was observed in control plots (Figure 3). Thus, shading affected both the timing and the dynamics of senescence, and it affected both aspects of senescence more in ‘Piznair’, which was also more strongly affected by shading in its sink potential.

**Figure 3.**
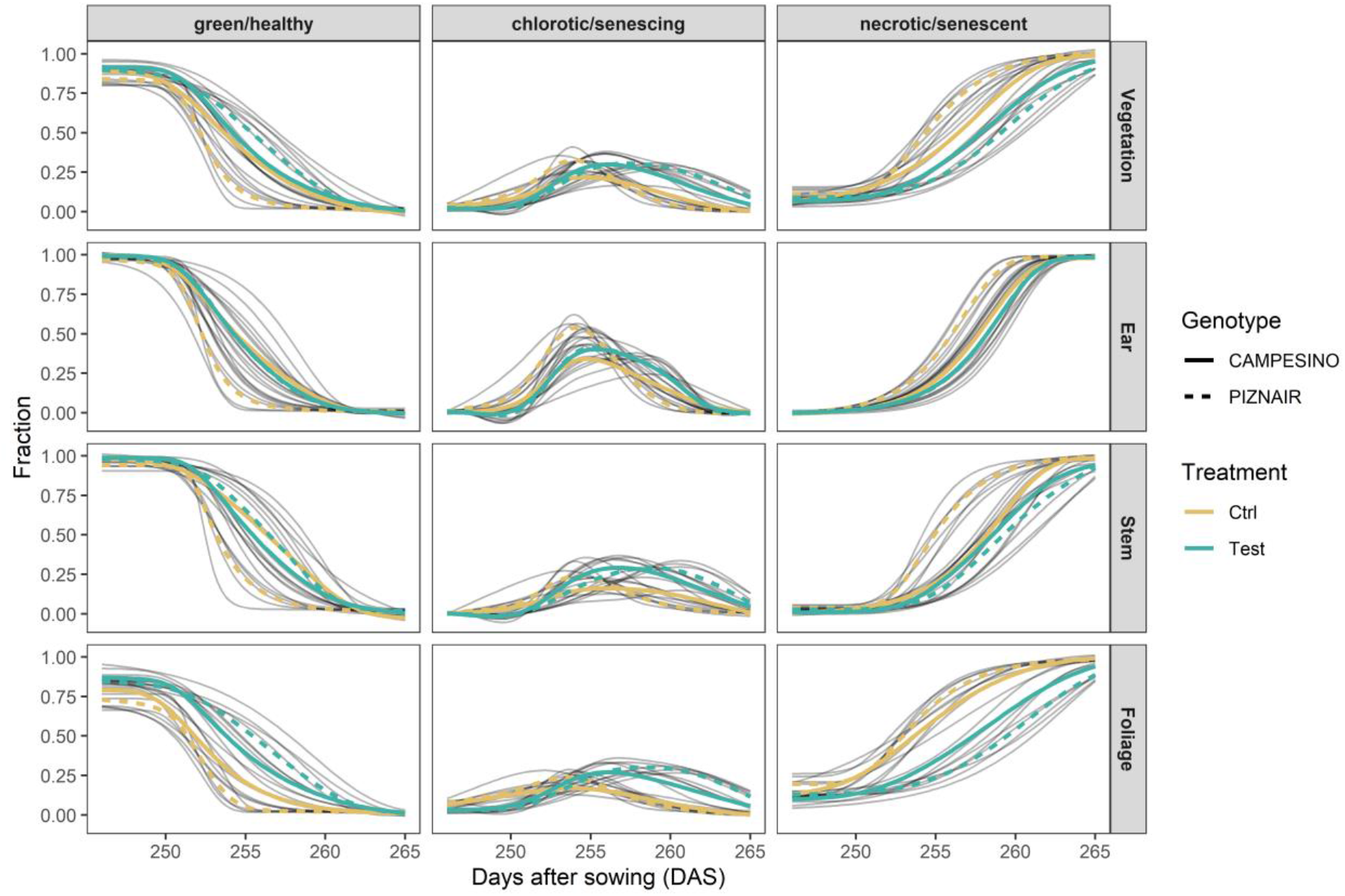
Relative contribution of green, chlorotic, and senescent tissue at organ level (total vegetation, ears, stems, and leaves) and their evolution over time between approximately 20 d after heading (21 June 2023) and physiological maturity (10 July 2023). Gray curves represent 4-parameter Gompertz model fits or P-spline fits to 16 data points for each experimental plot. Colored lines represent means for each treatment-by-genotype combination.

**Figure 4.**
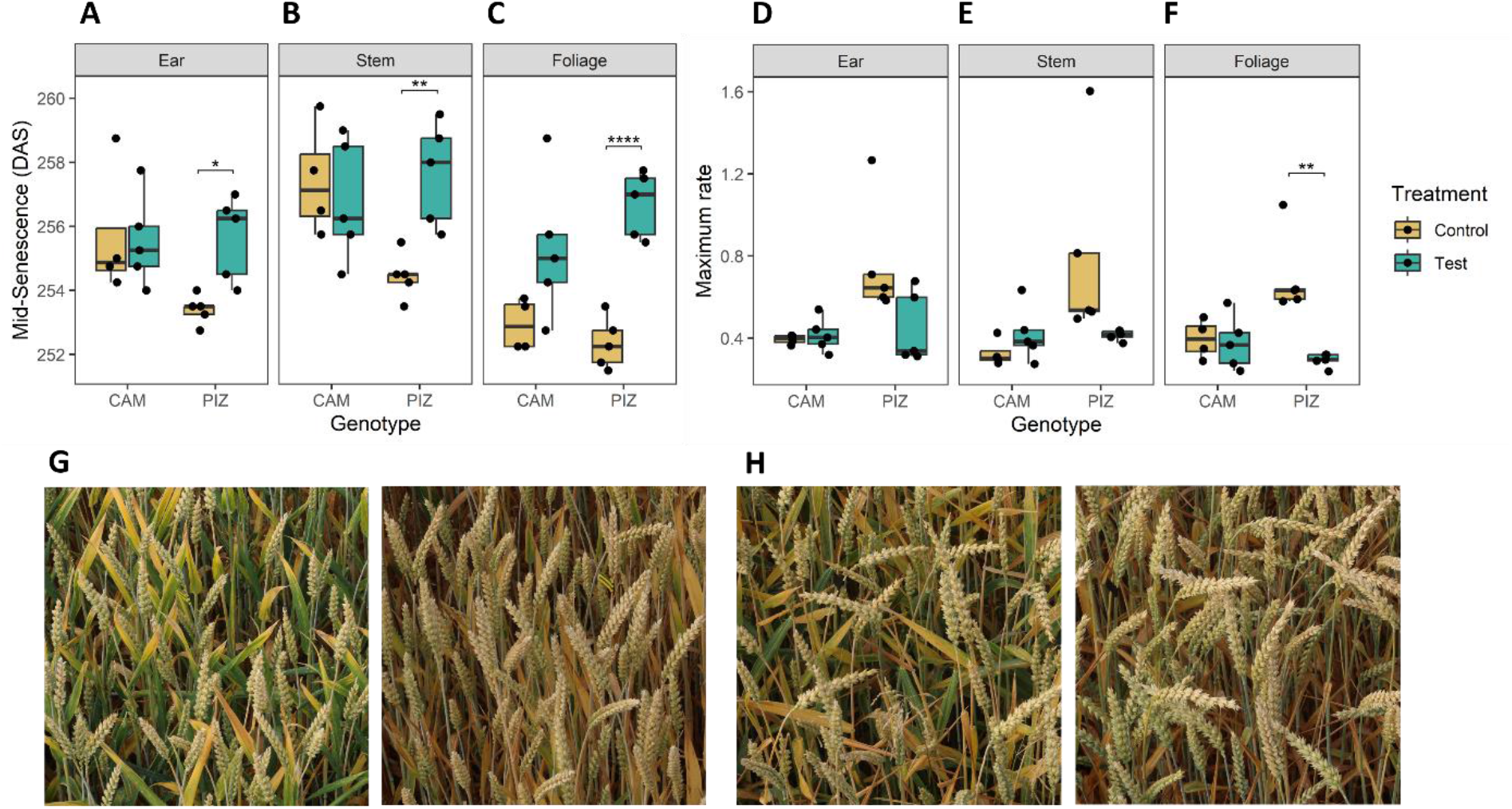
Effects of canopy shading on senescence dynamics. **(A)** Effects of shading on the midpoint of senescence observed for ears, **(B)** stems, **(C)** foliage; **(D)** Effects of shading on the maximum rate of senescence observed for ears, **(E)** stems, **(F)** foliage; **(G)** Example images of neighboring plots with shading (left) and without shading (right) for cultivar ‘Piznair’, **(H)** for cultivar ‘Campesino’. Images in (G) and (H) were acquired on 1 July 2023, i.e., at around the midpoint of senescence (256 DAS).

Overall, measurements of gas exchange and photosynthesis showed a high variation across measured leaves (raw data not shown), despite very stable weather conditions during the relevant period. Nonetheless, taken together the measurements indicated a trend towards reduced stomatal conductance and reduced photosynthetic activity in shaded plots compared to the controls (Figure 5), although these trends were not statistically significant for either of the tested cultivars. Interestingly, however, the stronger effects of shading on ‘Piznair’ compared to ‘Campesino’ as described above were by trend visible also in these measurements, with the only exception being the measurement of stomatal conductance at 245 DAS (20 June 2023), which represented the last measurement before the onset of visually observable senescence (cf. Figure 3). Across all measurement dates, the effects of shading were more pronounced in measurements of photosynthesis than in measurements of stomatal conductance for both tested cultivars (Figure 5). The Fq’/Fm’ showed the typical decrease with increasing PPFR which was more pronounced in ‘Piznair’ than in ‘Campesino’ (Figure 5C). In agreement with the photosynthesis measurements, shading slightly decreased Fq’/Fm’ compared to the control, again more pronounced in ‘Piznair’ (Figure 6D).

**Figure 5.**
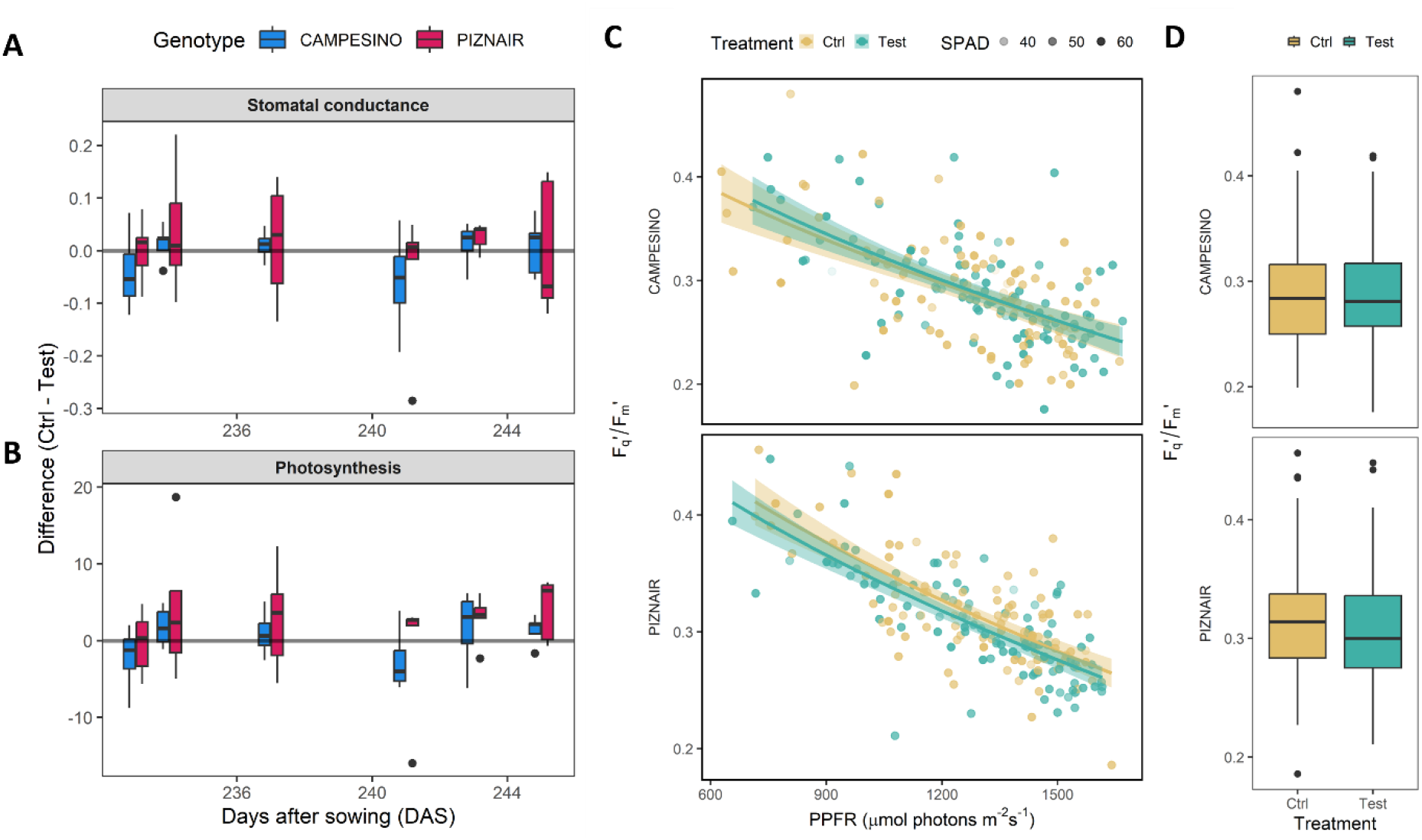
(A) Differences in stomatal conductance between members of pairs of directly adjacent plots, of which one each was exposed to pre-anthesis canopy shading (“Test”), and one was left untreated (“Control”). Positive values indicate higher stomatal conductance and photosynthetic rates in untreated control plots than in shaded test plots; **(B)** Differences in photosynthesis as determined via gas exchange. **(C)** Fq’/Fm’ in response to increased photosynthetic photon flux rate (PPFR). Solid lines represent fitted values after square root transformation of PPFR. Relative chlorophyll (SPAD) is indicated by the transparency of the points. **(D)** Boxplot of Fq’/Fm’ for both genotypes and treatments. Data for six measurement days between June 1 and June 14 (DAS 226 to 239) was pooled.

**Figure 6.**
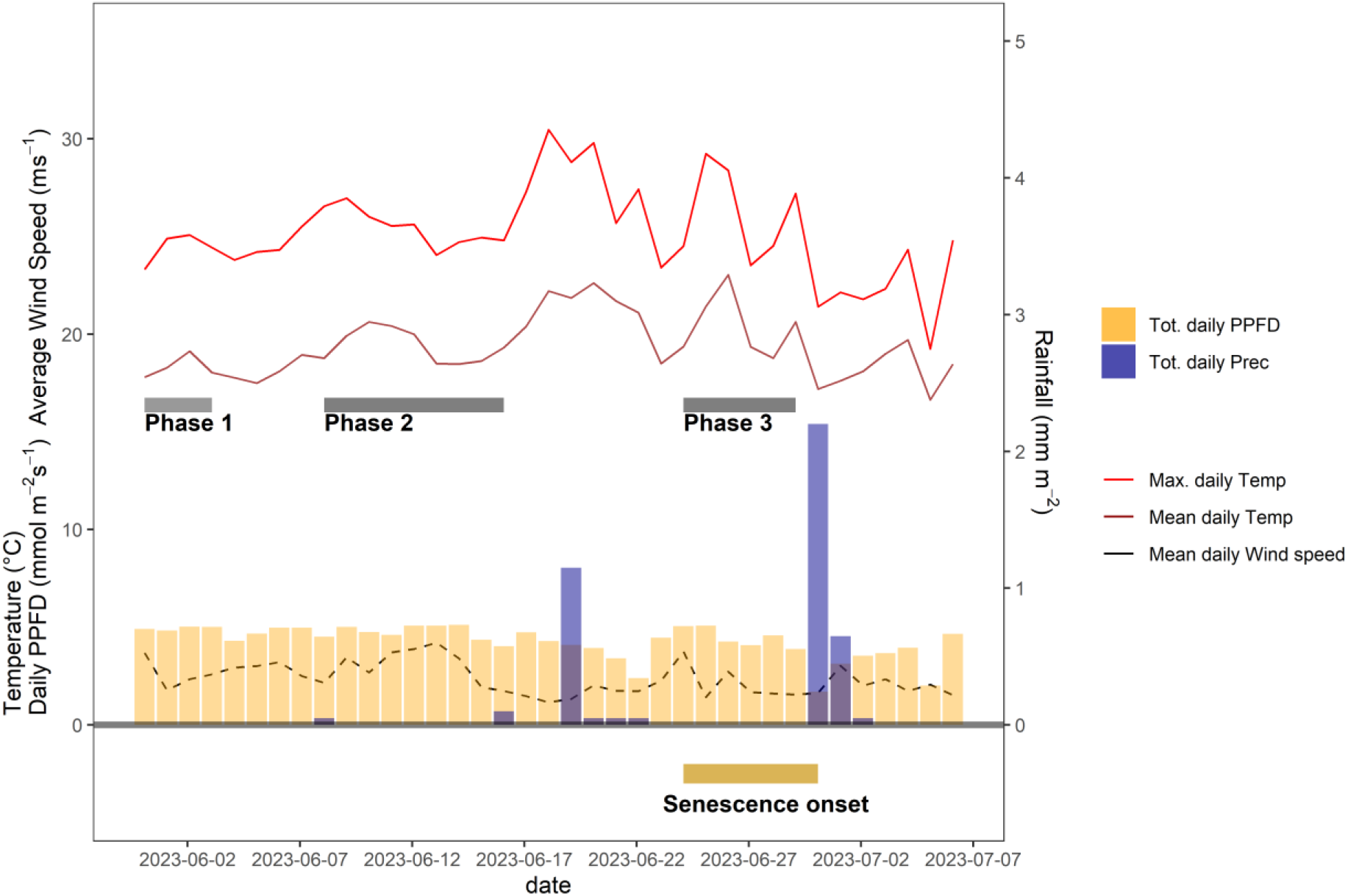
Daily average and maximum air temperature, daily rainfall, average daily wind speed, and daily incident light throughout the grain filling period of 2023. Except for rainfall, daily values were calculated based on data recorded between 10 a.m. and 4 p.m. each day. Dark grey bars indicate different phases during which distinct but temporally stable patterns in CT were observed. The dark yellow bar at the bottom of the plot indicates the time span during which plots started to senesce.

### Pre-anthesis canopy shading was associated with increasingly higher canopy temperature during early grain filling

CT values exhibited significant short-term variation, especially during early afternoons. Recurring rapid increases and decreases, typically in the range of 1-2 degrees, were observed within a timespan of a few minutes (cf. Supplementary Figure S3). Neighboring plots generally showed very similar temporal patterns in CT (Supplementary Figure S3), suggesting that this variation over time resulted from short-term changes in local meteorological conditions.

After a particularly wet spring season, early grain filling coincided with a period of very stable weather conditions, characterized by long periods of clear sky conditions during the day but moderate day-time temperatures (Figure 6). Significant rainfall was registered only on 19 June 2023, a few days before the onset of visually discernable senescence (Figure 6).

To isolate the effect of the treatment from effects related to genotype morphology, short-term changes in meteorological conditions, and spatial field heterogeneity, we calculated differences in CT between neighboring plots that had received either of the treatments, at the genotype level. When considering the entire measurement period (i.e., the period between flowering and harvest), contrasting effects of the treatment were observed (Figure 7). However, effects showed marked temporal continuity, allowing to distinguish three different phases (cf. Figure 6): In an initial Phase following the removal of the shading nets (Phase 1), shaded plots were consistently cooler during the day (Figure 7, top row). The difference between AUC_CT_ across the treatments were highly significant (p <0.01; Figure 8, top row), but this difference progressively disappeared within the first 4-5 d. During the following days (Phase 2), the initially observed trend was gradually inverted, with shaded plots increasingly characterized by higher CT than control plots (between 9 June 2023 and 16 June 2023; Figure 7, panel rows 2 and 3). This observed difference in AUC_CT_ was statistically significant for ‘Piznair’, but not for ‘Campesino’. When data for both genotypes was pooled, differences were at the significance threshold on most measurement dates (Figure 8). The trend observed during Phase 1 was therefore continued during Phase 2. The gradual increase in CT during Phase 1 and Phase 2 in shaded relative to control plots becomes more obvious when comparing AUC_CT_ across the entire duration (Figure 9). This also reveals that the relative increase appeared to be stronger for ‘Piznair’ than for ‘Campesino’. Finally, a third phase (Phase 3) with stable patterns across multiple days was observed between 23 June and 29 June, when shaded test plots were again warmer than control plots (bottom rows of Figures 7 and 8). This phase coincided with the onset of canopy senescence, which occurred earlier in control plots than in shaded test plots. Throughout the entire measurement period, differences between shaded and control plots did not exceed 1°C but were more often in the order of a few tens of degrees (Figure 7). Throughout the period, no sizable differences between differentially treated plots were observed during the night for either of the tested genotypes. Similarly, no differences in CT between shaded and control plots were apparent during the final days before maturity.

**Figure 7.**
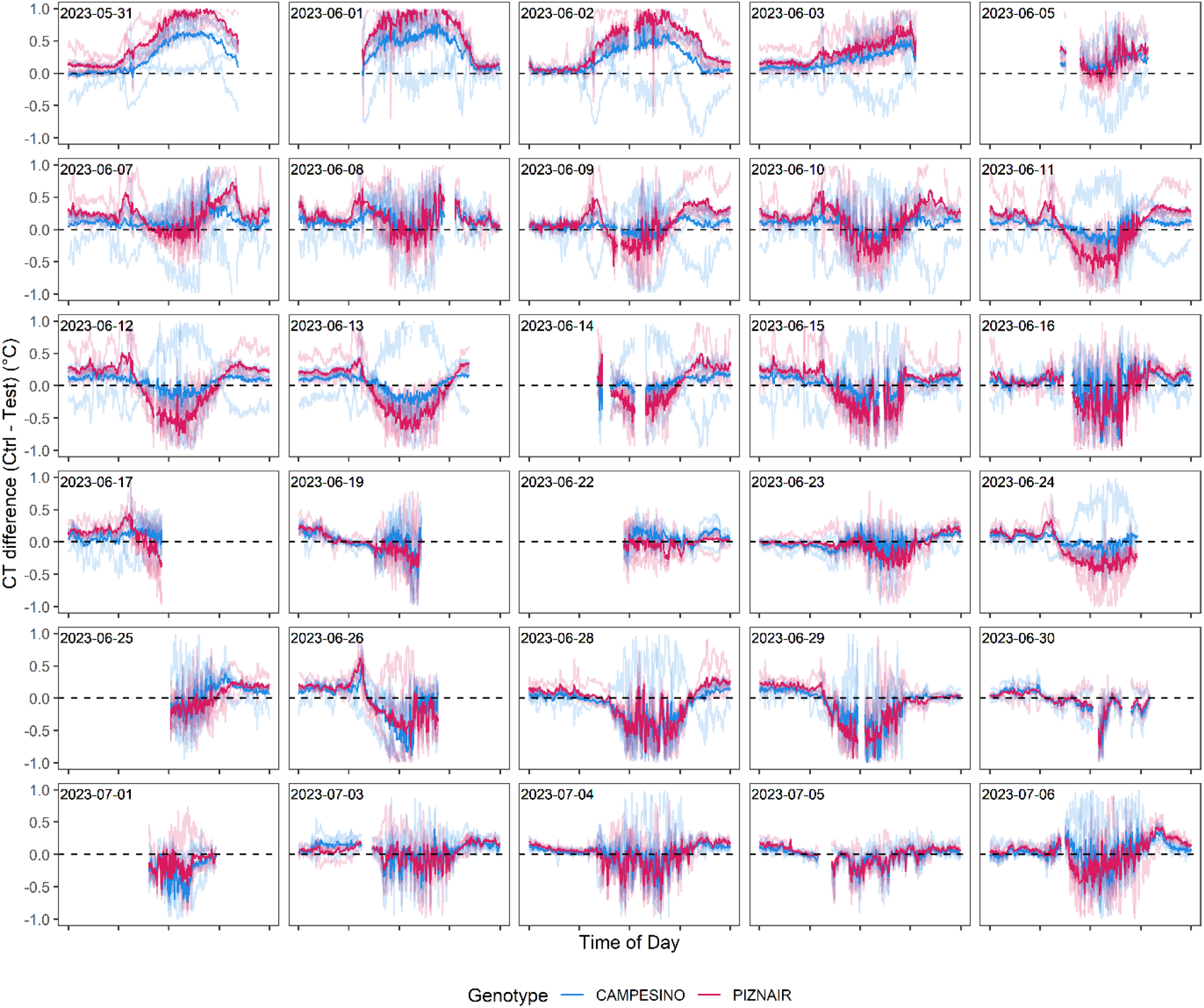
Differences in canopy temperature (CT) between pairs of plots with pre-anthesis shading (“Test”) and without pre-anthesis shading (“Ctrl”). Negative values indicate higher CT in test plots than in adjacent control plots, and vice-versa. Solid lines represent genotype averages across five replicates, transparent lines represent differences between individual pairs of neighboring plots. Measurement dates with little or no data available for the period between 10 a.m. and 6 p.m. are not included.

**Figure 8.**
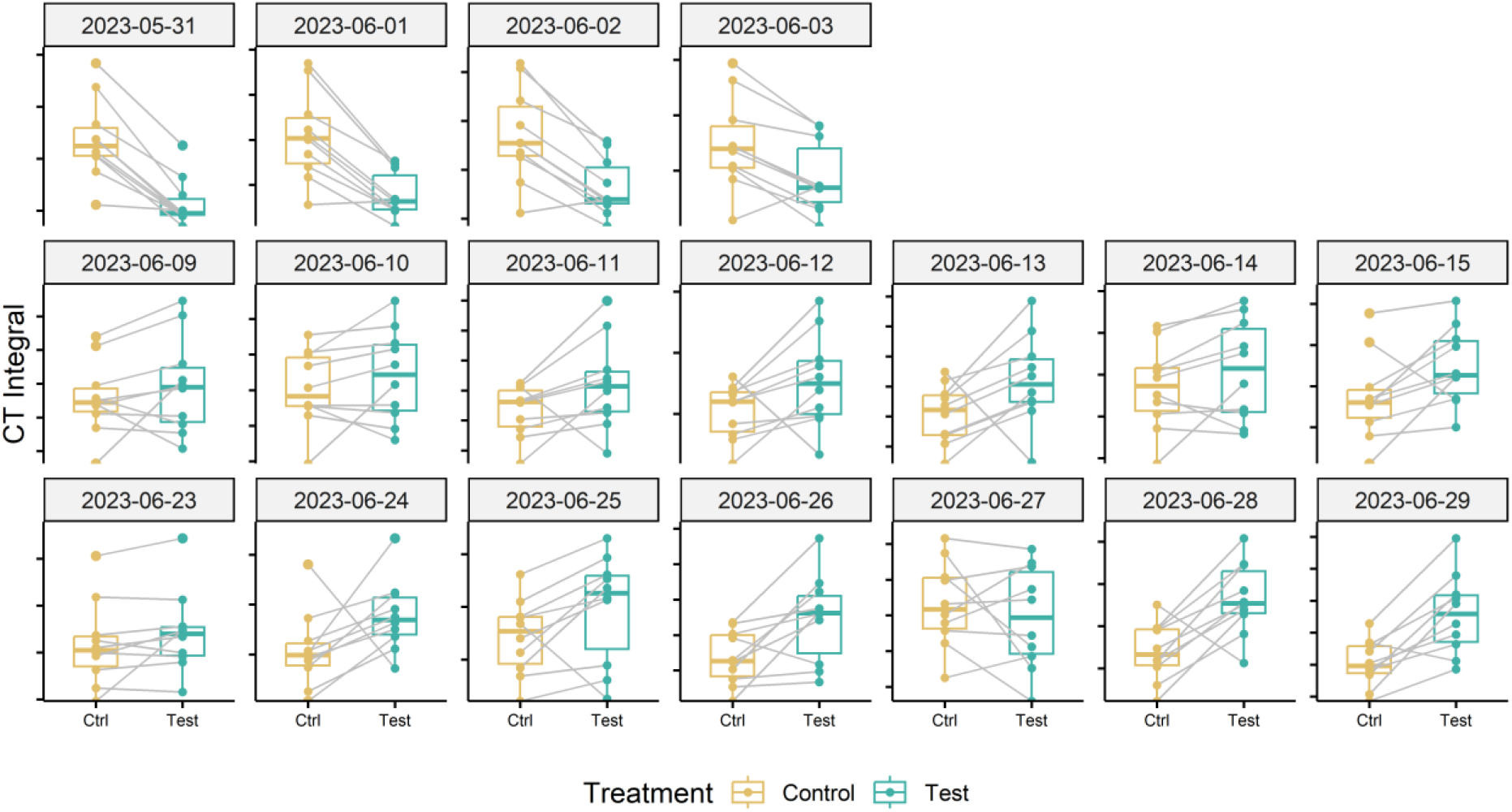
Effect of the pre-anthesis canopy shading treatment on the daily integral under the canopy temperature curves between 10 a.m. and 4 p.m. Data for measurement dates corresponding to three different phases of interest with stable environmental conditions are shown. Data for the two tested cultivars were pooled. Numbers above the black brackets indicate p-values of the paired sample t-test (n = 10), gray lines connect paired samples (i.e., neighboring plots of the same genotype having received the shading or control treatment).

## Discussion

Canopy temperature is often interpreted as representing leaf activity traits, particularly stomatal conductance, and corresponding associations have been found by many researchers (e.g., Fischer et al., 1998; Rebetzke et al., 2013). However, it is well known that CT measurements are strongly affected by a multitude of other factors as well (Anderegg et al., 2021; Deery and Jones, 2021; Prashar and Jones, 2014). Depending on the objective of CT measurements, the influence of confounding variables may or may not be problematic. For example, if reasonably strong correlations between CT and yield are consistently observed under certain well-defined growing conditions at well-defined stages of crop development (e.g., because CT may provide a fast measure of overall vigor under stressful conditions at that time), then an exact understanding of the underlying functional relationship is not necessary to use it as a trait to select for higher-yielding genotypes under these conditions (see Li et al., 2019; Reynolds et al., 1994; Thapa et al., 2018 for some examples). In contrast, if the objective of CT measurements is to complement other secondary traits (such as plant height and biomass) to enable a knowledge-based physiological selection, then a better understanding of the variables determining CT is indispensable. Only then will it be possible to estimate the feasibility of specific trait assessments, develop appropriate measurement protocols and define the scope of application. Here, our objective is of the latter sort: We aim to quantify a specific secondary trait (leaf activity traits during the stay-green phase of wheat), and we require this measurement to be *complementary* to (not indicative of) traits such as plant height, above-ground biomass, or phenology, for which more direct and precise high throughput assessment protocols are already available.

To better understand the potential and limitations of CT measurements in this context, we therefore performed an experiment that isolated the effect of variation in the trait of interest on CT from variation in CT caused by such confounding and possibly co-varying traits. Below, we first discuss the experimental approach underlying this feasibility assessment. Then, we interpret and contextualize observed effects and effect sizes, and discuss implications for an optimized use of CT measurements to characterize stay-green canopies.

### Pre-anthesis canopy shading to manipulate sink-source balances and create variation in stay-green functionality

We applied pre-anthesis canopy shading during rapid spike growth (but significantly after full flag leaf emergence) with the aim of creating variation in sink strength with minimum side-effects on the size and structure of the canopy. Given the importance of the sink-source balance in the regulation of senescence, shading was expected to result in dysfunctional stay-green *via* a reduced and delayed demand for remobilized assimilates and N.

Large effects of pre-anthesis shading on potential grain yield have been reported in several studies (e.g., Fischer, 1985, 1975; Savin and Slafer, 1991). Yield losses result primarily from a reduced number of kernels per spikelet (Slafer et al., 1994), and an increased number of rudimentary basal spikelets is often observed (Backhaus et al., 2023; Slafer et al., 1994), which appears to be the result of complete floret abortion in basal spikelets as a consequence of their delayed development (Backhaus et al., 2023). Though we did not perform detailed quantitative in-field assessments of spike fertility, it was obvious from ear volume measurements (Figure 2d) and from examinations of organ contribution to vegetation sceneries (Supplementary Figure S2) that shading drastically affected spike fertility. Specifically, there was an obvious increase in the number of rudimentary basal spikelets and completely aborted apical spikelets (Figure 2E; Figure 4D). Interestingly, eye-ball assessments in the field and re-examination of ears in images suggested that shading affected apical spikelets more than basal spikelets, especially in ‘Piznair’, which would be in contrast with common observation (Backhaus et al., 2023; Slafer et al., 1994; Stockman et al., 1983).

Because of the temporal overlap between rapid spike growth and elongation of the last internode (peduncle), canopy shading affected both traits (Figure 2). Peduncles are important as storage location for water-soluble carbohydrates, and canopy shading affects WSC accumulation within a few days of application (Stockman et al., 1983). Therefore, source strength must also have been affected by the shading treatment. However, this regarded primarily reserve and biomass accumulation of the peduncle and spike during the restricted period of shading, whereas photosynthetically active leaf area should not have been affected (Supplementary Figure S2). Given the size of the effects on sink potential, it seems highly likely that shading affected sink strength far more than source capacity, resulting in an increased source:sink ratio as compared to the untreated controls for both assimilates and Nitrogen. This is additionally supported by the increase in thousand kernel weight in ‘Campesino’ and the increase in grain protein concentration in response to shading in both tested genotypes. The lack of a sizable increase in TKW in ‘Piznair’ is likely a result of grain size limitations (i.e., the maximum capacity of grains to absorb assimilates, that is determined already at an early stage of grain filling (Borrás et al., 2004; Brocklehurst, 1977)) than the lack of assimilate supply, given that GPC is drastically increased despite its already high levels under control conditions.

The dynamics of senescence were strongly affected by shading, with magnitude and direction of the observed changes in good agreement with an interpretation of the shading treatment as a cause of dysfunctional stay-green mediated by source:sink imbalance. In apparently sink-limited ‘Campesino’, shading reduced grain number, but this was partly compensated for by a sizeable increase in TKW and in GPC (Figure 2E, 2F), which helped maintain a high sink demand for assimilates and remobilized Nitrogen. Consequently, only relatively small differences between treatments in the timing and dynamics of senescence were observed in this cultivar, and only for leaves (Figure 2). In contrast, ‘Piznair’ was unable to compensate *via* increases in TKW, and sink demand for assimilates and Nitrogen was therefore more strongly reduced. Accordingly, the delay in the onset and progression of senescence was accentuated in this cultivar, and the delay was also apparent in ears and stems. These patterns therefore confirm the importance of sink:source balances in the regulation of senescence under stress-free conditions and strongly suggest that pre-anthesis canopy shading caused dysfunctional stay-green.

In contrast to the obvious agronomic and phenological effects of shading, direct measurements of photosynthetic rate and stomatal conductance in the field were less conclusive, although direction and relative size of effects across the tested genotypes do seem to indicate that the expected decrease in photosynthetic rate and stomatal conductance did in fact occur (Figure 5). Reynolds et al. (2005) and Miralles and Slafer (1997) also suggested that sink strength should be a major factor determining post-anthesis growth, and a common observation has been that upon removal of source capacity during grain filling, stomatal conductance and photosynthetic rates are increased in compensation (Rawson et al., 1976; Richards, 1996); so the inverse should equally be true. In a similar experiment with zucchini, sink limitation did not significantly reduce photosynthetic rates, but nevertheless increased leaf temperature and had other notable leaf- and canopy level effects that were readily detectable using reflectance-based approaches (Burnett et al., 2021). Thus, sink-limitation may not necessarily have immediate and strong effects on leaf photosynthesis. Here, an additional plausible explanation for the lack of a similarly clear difference between treatments in stomatal conductance and photosynthesis may be the difficulty in upscaling spot measurements performed on single and randomly selected leaves to entire field canopies.

### Feasibility of stay-green functionality monitoring using high throughput canopy temperature measurements

CT differed significantly between the treatments in the initial phase immediately following the removal of the shading tents (Figure 7, Figure 8, top rows), and this effect cannot be well explained by expected differences in leaf activity traits resulting from the shading treatment. It seems more likely that these initial differences represented side-effects of the shading treatment on canopy structure and on the contribution of different organs to plot-level CT signals (Supplementary Figure S2). Spikes and peduncles were found to be consistently warmer than flag leaves at different stages of grain filling by Ayeneh et al. (2002); Similarly, Vicente et al. (2018) found durum wheat ears to be consistently warmer than leaves, and Fernandez-Gallego et al. (2019) even exploited this fact to obtain automatic ear counts from thermal images. A higher contribution of warmer ears to plot-level CT signals is therefore a likely reason for the observed initial differences (cf. Supplementary Figure S2). When considering the entire stay-green phase, there was an obvious gradual increase in CT of shaded canopies relative to unshaded controls over a period of almost three weeks. This increase appeared to be nearly constant during the early grain filling phase (until approximately 18 June) and was then interrupted by a period of less stable weather conditions (Figure 6); however, it quickly reestablished upon a return of more stable weather conditions shortly after. These basic observations are in full agreement with the indications based on all other measurements that shading introduced dysfunctional stay-green and it clearly suggests that this had a direct and measurable effect on CT. In particular, a gradually increasing contrast in CT between treatments is expected under the assumption of an increasingly downregulating effect of limited sink demand on grain filling rates and, consequently, leaf activity traits. Conversely, a more abrupt effect would be expected if the reduction in sink demand immediately triggered a proportional decrease in leaf activity. Unfortunately, to the best of our knowledge, sink-regulation of photosynthesis during grain filling in wheat (and thus the expected response of photosynthetic rates and stomatal conductance to reductions in sink strength) is poorly understood. Intuitively, it would seem more natural for this effect to build up gradually, since there should be no constraints to grain filling in an initial phase.

Despite the strong effects of the shading treatment on physiological and phenological traits of interest, observed differences in CT between the treatments were limited to less than 1°C. This is substantially less variation than we observed previously within a set of diverse genotypes measured during the same growth stages at the same site (Anderegg et al., 2021; Perich et al., 2020), where differences across measurement dates between coolest and hottest canopies ranged from 3.1°C to 11.8°C in raw data, and from 1.8°C to 6.8°C after spatial correction of the CT signals (reanalyzed from Anderegg et al., 2021; Perich et al., 2020). Limiting these analyses to Swiss elite breeding material resulted in a very similar picture. Similar ranges were observed across different years and time of day by Deery et al. (2019, see e.g., Figure 9 in their paper). If our interpretation of the initial differences in CT is correct, then that should be considered the baseline for the quantification of the effect of differences in stay-green functionality on CT, i.e., the total effect would amount to approximately 1.5°C and 0.8°C for ‘Piznair’ and ‘Campesino’, respectively (cf. Figure 7). While this would be a sizeable effect, it clearly must be considered the maximum expected effect in experiments without treatments. In a set of historical lines from CIMMYT spanning 26 years of breeding, reported a decrease in CT of approximately 0.6°C that could be associated with higher stomatal conductance (Fischer et al., 1998). Yet, this number may have been influenced by traits other than leaf activity, although above ground biomass was ruled out as a relevant factor in that study.

**Figure 9.**
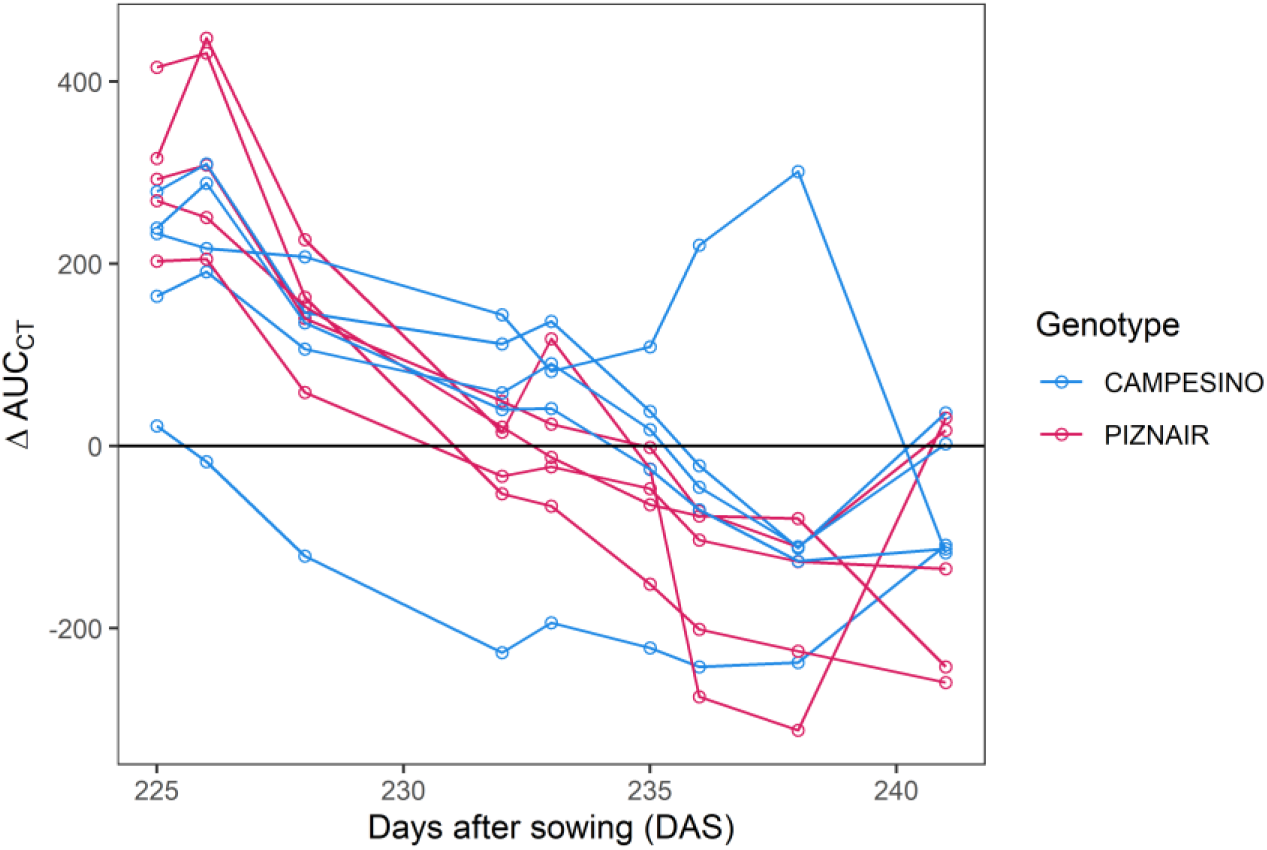
Differences between control plots and shaded plots in the area under the canopy temperature curve observed between 10 a.m. and 4 p.m. on each day (AUCCT). Data is shown for the early grain filling phase i.e., up until 20 June 2023, which corresponds to the phase preceding the onset of visually discernable senescence in the earliest maturing plots.

The gradual nature of the observed effects of sink limitation on CT during grain filling confirms the hypothesized advantage of a time-integrated analysis of individual measurements (Anderegg et al., 2021). We already observed a moderate to high heritability of plot-based temporal trends in CT under the conditions of the study site, but it was unclear to what extent they represented genotype-specific reactions to progressive soil drying or again an effect of confounding factors, such as heritable changes in canopy structure during the assessment period. Given the possibility of within-genotype comparisons in this study, these confounding effects could be excluded, underscoring the meaningfulness and advantage of time-integrated analysis of CT changes during periods of stable meteorological conditions.

## Conclusions

This study integrated gold standard physiological measurements and traditional experimental approaches with recently developed RGB- and thermal-image-based high throughput phenotyping protocols, with the aim of developing a better understanding of the sensitivity limits of remote-sensing-based phenotyping approaches in stay-green wheat canopies. Our results clearly indicate that differences in leaf activity stemming from differences in stay-green functionality translate into measurable differences in CT. Importantly, they appear to do so in the absence of co-variation in confounding traits (such as above ground biomass, green leaf area, or plant height), and in dense canopies with (near-)complete soil cover, as typically observed in high-yielding, stress-free environments. Our findings provide a strong basis for future uses of CT for a better characterization of the physiological status of stay-green wheat canopies during early grain filling. Modest effect sizes highlight the importance of restricting screenings to a limited range of morphological and phenological diversity, as already recommended in a similar context by Lopes and Reynolds (2010). Finally, gradually increasing effects of sink limitation on CT underscore the importance of frequent measurements and a time-integrated analysis.

## Author contributions

JA proposed and designed the experiment, performed in-field and post-harvest measurements, processed, analyzed, and interpreted the data, and wrote the original draft of the manuscript. AH contributed to the conceptualization and design of the study, assisted with field assessments, contributed to data interpretation, provided materials and resources, and reviewed and edited the manuscript. NK designed, maintained, mounted, and operated the multi-sensor pack that included the thermal camera, set up the ear volume estimation method, assisted with experiment planning, and reviewed and edited the manuscript. HA acquired funding for and designed the multi-sensor pack, and reviewed and edited the manuscript. OZ collected and processed ear volume data, and reviewed and edited the manuscript. BK contributed to in-field measurements, contributed to data interpretation, and reviewed and edited the manuscript. RZ assisted with deep learning model development, provided annotated data, provided technical support with computing infrastructure, and reviewed and edited the manuscript. AW contributed to the conceptualization of the study, contributed materials and resources, and reviewed and edited the manuscript.

## Funding

We acknowledge funding for the development and implementation of the multi-sensor pack within the Project CT-PhotoInd, funded under the SNF SPARK theme (Project Number CRSK-3_195591). The rest of the project was funded by ETH Zürich.

## Acknowledgements

We thank Simon Corrado and Brigitta Herzog (both ETH Crop Science) for crop husbandry and assistance with seed preparation and management; Seraina Wagner, Sophie Vuillemin, Tenzin Khampo (all ETH Plant Pathology) and Michelle Kremers (ETH Crop Science) for assistance with field measurements and annotations; Mohammad Hosein Fakhrosae (ETH Crop Science) for help with harvest trait determination; and Bruce McDonald (ETH Plant Pathology) for sharing resources and for assistance in project coordination and management.

## Notes

### Competing Interest Statement

The authors have declared no competing interest.

## References

1. Amani, I., Fischer, R.A., Reynolds, M.P., 1996. Canopy Temperature Depression Association with Yield of Irrigated Spring Wheat Cultivars in a Hot Climate. Journal of Agronomy and Crop Science 176, 119–129. 10.1111/j.1439-037X.1996.tb00454.x

2. Anderegg, J., Aasen, H., Perich, G., Roth, L., Walter, A., Hund, A., 2021. Temporal trends in canopy temperature and greenness are potential indicators of late-season drought avoidance and functional stay-green in wheat. Field Crops Research 274, 108311. 10.1016/j.fcr.2021.108311

3. Anderegg, J., Yu, K., Aasen, H., Walter, A., Liebisch, F., Hund, A., 2020. Spectral Vegetation Indices to Track Senescence Dynamics in Diverse Wheat Germplasm. Front. Plant Sci. 10. 10.3389/fpls.2019.01749

4. Anderegg, J., Zenkl, R., Walter, A., Hund, A., McDonald, B.A., 2023. Combining High-Resolution Imaging, Deep Learning, and Dynamic Modeling to Separate Disease and Senescence in Wheat Canopies. Plant Phenomics 5, 0053. 10.34133/plantphenomics.0053

5. Ayeneh, A., van Ginkel, M., Reynolds, M.P., Ammar, K., 2002. Comparison of leaf, spike, peduncle and canopy temperature depression in wheat under heat stress. Field Crops Research 79, 173–184. 10.1016/S0378-4290(02)00138-7

6. Backhaus, A.E., Griffiths, C., Vergara-Cruces, A., Simmonds, J., Lee, R., Morris, R.J., Uauy, C., 2023. Delayed development of basal spikelets in wheat explains their increased floret abortion and rudimentary nature. Journal of Experimental Botany 74, 5088–5103. 10.1093/jxb/erad233

7. Borrás, L., Slafer, G.A., Otegui, M.E., 2004. Seed dry weight response to source–sink manipulations in wheat, maize and soybean: a quantitative reappraisal. Field Crops Research 86, 131–146. 10.1016/j.fcr.2003.08.002

8. Brocklehurst, P.A., 1977. Factors controlling grain weight in wheat. Nature 266, 348–349. 10.1038/266348a0

9. Burnett, A.C., Serbin, S.P., Rogers, A., 2021. Source:sink imbalance detected with leaf- and canopy-level spectroscopy in a field-grown crop. Plant, Cell & Environment 44, 2466– 2479. 10.1111/pce.14056

10. Chapman, E.A., Orford, S., Lage, J., Griffiths, S., 2021. Capturing and Selecting Senescence Variation in Wheat. Front. Plant Sci. 12. 10.3389/fpls.2021.638738

11. Deery, D.M., Jones, H.G., 2021. Field Phenomics: Will It Enable Crop Improvement? Plant Phenomics 2021. 10.34133/2021/9871989

12. Deery, D.M., Rebetzke, G.J., Jimenez-Berni, J.A., Bovill, W.D., James, R.A., Condon, A.G., Furbank, R.T., Chapman, S.C., Fischer, R.A., 2019. Evaluation of the Phenotypic Repeatability of Canopy Temperature in Wheat Using Continuous-Terrestrial and Airborne Measurements. Front. Plant Sci. 10. 10.3389/fpls.2019.00875

13. Deery, D.M., Rebetzke, G.J., Jimenez-Berni, J.A., James, R.A., Condon, A.G., Bovill, W.D., Hutchinson, P., Scarrow, J., Davy, R., Furbank, R.T., 2016. Methodology for High-Throughput Field Phenotyping of Canopy Temperature Using Airborne Thermography. Front. Plant Sci. 7. 10.3389/fpls.2016.01808

14. Fernandez-Gallego, J.A., Kefauver, S.C., Vatter, T., Aparicio Gutiérrez, N., Nieto-Taladriz, M.T., Araus, J.L., 2019. Low-cost assessment of grain yield in durum wheat using RGB images. European Journal of Agronomy 105, 146–156. 10.1016/j.eja.2019.02.007

15. Fischer, R.A., 1985. Number of kernels in wheat crops and the influence of solar radiation and temperature. The Journal of Agricultural Science 105, 447–461. 10.1017/S0021859600056495

16. Fischer, R.A., 1975. Yield Potential in a Dwarf Spring Wheat and the Effect of Shading1. Crop Science 15, cropsci1975.0011183X001500050002x. 10.2135/cropsci1975.0011183X001500050002x

17. Fischer, R.A., Rees, D., Sayre, K.D., Lu, Z.-M., Condon, A.G., Saavedra, A.L., 1998. Wheat Yield Progress Associated with Higher Stomatal Conductance and Photosynthetic Rate, and Cooler Canopies. Crop Science 38, 1467. 10.2135/cropsci1998.0011183X003800060011x

18. Gregersen, P.L., Culetic, A., Boschian, L., Krupinska, K., 2013. Plant senescence and crop productivity. Plant Mol Biol 82, 603–622. 10.1007/s11103-013-0013-8

19. Gregersen, P.L., Holm, P.B., Krupinska, K., 2008. Leaf senescence and nutrient remobilisation in barley and wheat. Plant Biology 10, 37–49. 10.1111/j.1438-8677.2008.00114.x

20. Grieder, C., Hund, A., Walter, A., 2015. Image based phenotyping during winter: a powerful tool to assess wheat genetic variation in growth response to temperature. Functional Plant Biol. 42, 387–396. 10.1071/FP14226

21. He, K., Zhang, X., Ren, S., Sun, J., 2016. Deep Residual Learning for Image Recognition, in: 2016 IEEE Conference on Computer Vision and Pattern Recognition (CVPR). Presented at the 2016 IEEE Conference on Computer Vision and Pattern Recognition (CVPR), IEEE, Las Vegas, NV, USA, pp. 770–778. 10.1109/CVPR.2016.90

22. Jiang, G.H., He, Y.Q., Xu, C.G., Li, X.H., Zhang, Q., 2004. The genetic basis of stay-green in rice analyzed in a population of doubled haploid lines derived from an indica by japonica cross. Theor Appl Genet 108, 688–698. 10.1007/s00122-003-1465-z

23. Jones, H.G., Vaughan, R.A., 2011. Remote sensing of vegetation: principles, techniques and applications. By Hamlyn G. Jones and Robin A Vaughan. Journal of Vegetation Science 22, 1151–1153. 10.1111/j.1654-1103.2011.01319.x

24. Keller, B., Soto, J., Steier, A., Portilla-Benavides, A.E., Raatz, B., Studer, B., Walter, A., Muller, O., Urban, M.O., 2023. Linking photosynthesis and yield reveals a strategy to improve light use efficiency in a climbing bean breeding population. Journal of Experimental Botany erad416. 10.1093/jxb/erad416

25. Kichey, T., Hirel, B., Heumez, E., Dubois, F., Le Gouis, J., 2007. In winter wheat (Triticum aestivum L.), post-anthesis nitrogen uptake and remobilisation to the grain correlates with agronomic traits and nitrogen physiological markers. Field Crops Research 102, 22–32. 10.1016/j.fcr.2007.01.002

26. Kirchgessner, N., Liebisch, F., Yu, K., Pfeifer, J., Friedli, M., Hund, A., Walter, A., 2017. The ETH field phenotyping platform FIP: a cable-suspended multi-sensor system. Functional Plant Biol. 44, 154–168. 10.1071/FP16165

27. Kuhlgert, S., Austic, G., Zegarac, R., Osei-Bonsu, I., Hoh, D., Chilvers, M.I., Roth, M.G., Bi, K., TerAvest, D., Weebadde, P., Kramer, D.M., 2016. MultispeQ Beta: a tool for large-scale plant phenotyping connected to the open PhotosynQ network. Royal Society Open Science 3, 160592. 10.1098/rsos.160592

28. Lenth, R., Buerkner, P., Herve, M., Love, J., Riebl, H., Singmann, H., 2020. emmeans: Estimated Marginal Means, aka Least-Squares Means.

29. Li, X., Ingvordsen, C.H., Weiss, M., Rebetzke, G.J., Condon, A.G., James, R.A., Richards, R.A., 2019. Deeper roots associated with cooler canopies, higher normalized difference vegetation index, and greater yield in three wheat populations grown on stored soil water. J Exp Bot 70, 4963–4974. 10.1093/jxb/erz232

30. Lopes, M.S., Reynolds, M.P., 2010. Partitioning of assimilates to deeper roots is associated with cooler canopies and increased yield under drought in wheat. Functional Plant Biol. 37, 147–156. 10.1071/FP09121

31. Miralles, D.J., Slafer, G.A., 1997. Radiation interception and radiation use efficiency of near-isogenic wheat lines with different height. Euphytica 97, 201–208. 10.1023/A:1003061706059

32. Naruoka, Y., Sherman, J.D., Lanning, S.P., Blake, N.K., Martin, J.M., Talbert, L.E., 2012. Genetic Analysis of Green Leaf Duration in Spring Wheat. Crop Science 52, 99. 10.2135/cropsci2011.05.0269

33. Pask, A., Pietragalla, J., Mullan, D., Reynolds, M.P., 2012. Physiological breeding II : a field guide to wheat phenotyping iv, 132 pages.

34. Perich, G., Hund, A., Anderegg, J., Roth, L., Boer, M.P., Walter, A., Liebisch, F., Aasen, H., 2020. Assessment of Multi-Image Unmanned Aerial Vehicle Based High-Throughput Field Phenotyping of Canopy Temperature. Front. Plant Sci. 11. 10.3389/fpls.2020.00150

35. Pinheiro, J., Bates, D., DebRoy, S., Sarkar, D., 2021. nlme: Linear and Nonlinear Mixed Effects Models.

36. Prashar, A., Jones, H., 2014. Infra-Red Thermography as a High-Throughput Tool for Field Phenotyping. Agronomy 4, 397–417. 10.3390/agronomy4030397

37. R Core Team, 2018. R: A language and environment for statistical computing. R Foundation for Statistical Computing, Vienna, Austria. 2012. URL http://www.R-project.org.

38. Rajcan, I., Tollenaar, M., 1999. Source : sink ratio and leaf senescence in maize:: II. Nitrogen metabolism during grain filling. Field Crops Research 60, 255–265. 10.1016/S0378-4290(98)00143-9

39. Rawson, H.M., Gifford, R.M., Bremner, P.M., 1976. Carbon dioxide exchange in relation to sink demand in wheat. Planta 132, 19–23. 10.1007/BF00390326

40. Rebetzke, G.J., Rattey, A.R., Farquhar, G.D., Richards, R.A., Condon, A. (Tony) G., 2013. Genomic regions for canopy temperature and their genetic association with stomatal conductance and grain yield in wheat. Functional Plant Biol. 40, 14–33. 10.1071/FP12184

41. Reynolds, M.P., Balota, M., Delgado, M.I.B., Amani, I., Fischer, R.A., 1994. Physiological and Morphological Traits Associated With Spring Wheat Yield Under Hot, Irrigated Conditions. Functional Plant Biol. 21, 717–730. 10.1071/pp9940717

42. Reynolds, M.P., Pellegrineschi, A., Skovmand, B., 2005. Sink-limitation to yield and biomass: a summary of some investigations in spring wheat. Annals of Applied Biology 146, 39–49. 10.1111/j.1744-7348.2005.03100.x

43. Richards, R.A., 1996. Increasing yield potential in wheat - source and sink limitations., in: Increasing Yield Potential in Wheat: Breaking the Barriers.

44. Roche, D., 2015. Stomatal Conductance Is Essential for Higher Yield Potential of C3 Crops. Critical Reviews in Plant Sciences 34, 429–453. 10.1080/07352689.2015.1023677

45. Savin, R., Slafer, G.A., 1991. Shading effects on the yield of an Argentinian wheat cultivar. The Journal of Agricultural Science 116, 1–7. 10.1017/S0021859600076085

46. Slafer, G.A., Calderini, D.F., Miralles, D.J., Dreccer, M.F., 1994. Preanthesis shading effects on the number of grains of three bread wheat cultivars of different potential number of grains. Field Crops Research 36, 31–39. 10.1016/0378-4290(94)90050-7

47. Stockman, Y.M., Fischer, R.A., Brittain, E.G., 1983. Assimilate Supply and Floret Development Within the Spike of Wheat (Triticum aestivum L.). Functional Plant Biol. 10, 585–594. 10.1071/pp9830585

48. Strebel, S., Levy Häner, L., Mattin, M., Schaad, N., Morisoli, R., Watroba, M., Girard, M., Courvoisier, N., Berberat, J., Grandgirard, R., Graf, B., Streit, M., Weisflog, T., 2022. Liste der empfohlenen Getreidesorten für die Ernte 2023. Agroscope Transfer 443.

49. Thapa, S., Jessup, K.E., Pradhan, G.P., Rudd, J.C., Liu, S., Mahan, J.R., Devkota, R.N., Baker, J.A., Xue, Q., 2018. Canopy temperature depression at grain filling correlates to winter wheat yield in the U.S. Southern High Plains. Field Crops Research 217, 11–19. 10.1016/j.fcr.2017.12.005

50. Thomas, H., Ougham, H., 2014. The stay-green trait. J Exp Bot 65, 3889–3900. 10.1093/jxb/eru037

51. Treier, S., Roth, L., Hund, A., Kirchgessner, N., Aasen, H., Walter, A., Herrera, J.M., 2023. Digital lean phenotyping methods in the context of wheat variety testing – the cases of canopy temperature and phenology (preprint). Preprints. 10.22541/essoar.169870265.59272601/v1

52. Triboi, E., Triboi-Blondel, A.-M., 2002. Productivity and grain or seed composition: a new approach to an old problem—invited paper. European Journal of Agronomy 16, 163–186. 10.1016/S1161-0301(01)00146-0

53. Uauy, C., Brevis, J.C., Dubcovsky, J., 2006. The high grain protein content gene Gpc-B1 accelerates senescence and has pleiotropic effects on protein content in wheat. J Exp Bot 57, 2785–2794. 10.1093/jxb/erl047

54. van Oosterom, E.J., Chapman, S.C., Borrell, A.K., Broad, I.J., Hammer, G.L., 2010. Functional dynamics of the nitrogen balance of sorghum. II. Grain filling period. Field Crops Research 115, 29–38. 10.1016/j.fcr.2009.09.019

55. Vicente, R., Vergara-Díaz, O., Medina, S., Chairi, F., Kefauver, S.C., Bort, J., Serret, M.D., Aparicio, N., Araus, J.L., 2018. Durum wheat ears perform better than the flag leaves under water stress: Gene expression and physiological evidence. Environmental and Experimental Botany 153, 271–285. 10.1016/j.envexpbot.2018.06.004

56. Xie, Q., Mayes, S., Sparkes, D.L., 2016. Early anthesis and delayed but fast leaf senescence contribute to individual grain dry matter and water accumulation in wheat. Field Crops Research 187, 24–34. 10.1016/j.fcr.2015.12.009

57. Yang, J., Zhang, J., 2006. Grain filling of cereals under soil drying. New Phytologist 169, 223–236. 10.1111/j.1469-8137.2005.01597.x

