## Supplementary Figures for "Thermal imaging can reveal variation in stay-green functionality of wheat canopies under temperate conditions"

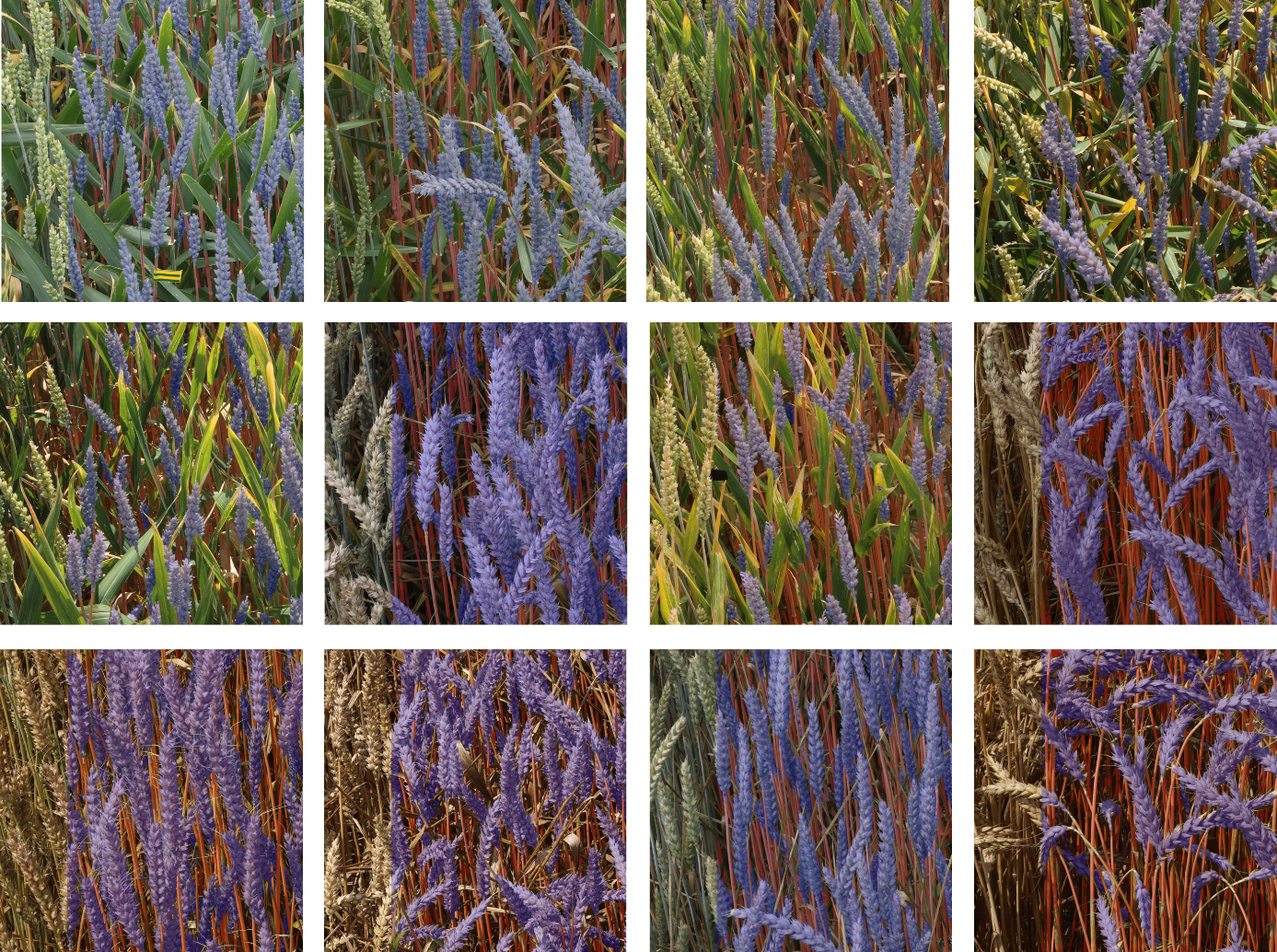

Supplementary Figure S 1 Inference on randomly selected images using the developed ear and stem segmentation model for off-nadir images. These segmentations were used to track organ-level senescence dynamics by means of organ-level color analysis throughout the grain filling phase.

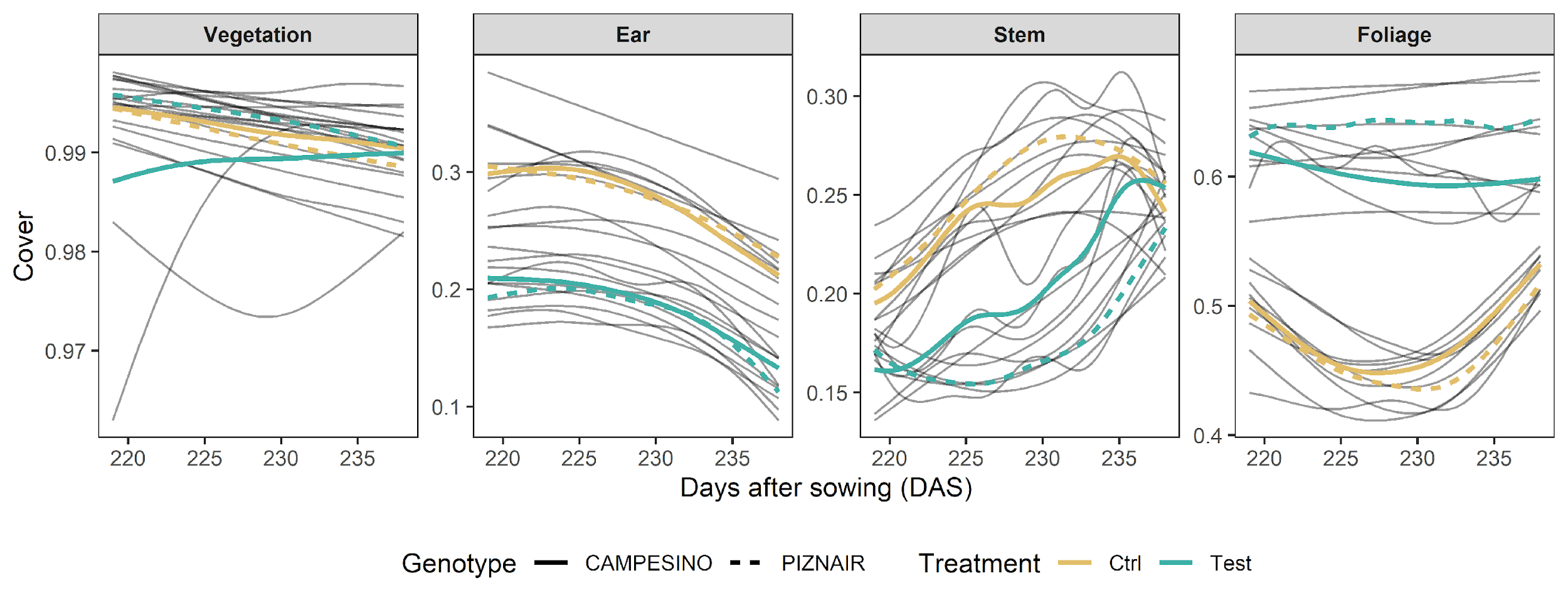

*Supplementary Figure S 2 Contribution of different organs to canopy sceneries as observed from an off-nadir perspective throughout the assessment period, as dependent on genotype and treatment. Individual black lines represent smoothed trends over time for individual plots (five repetitions per genotype- treatment combination). Note the variable scale of the y- axis for better visibility of treatment and genotype differences, where existent.*

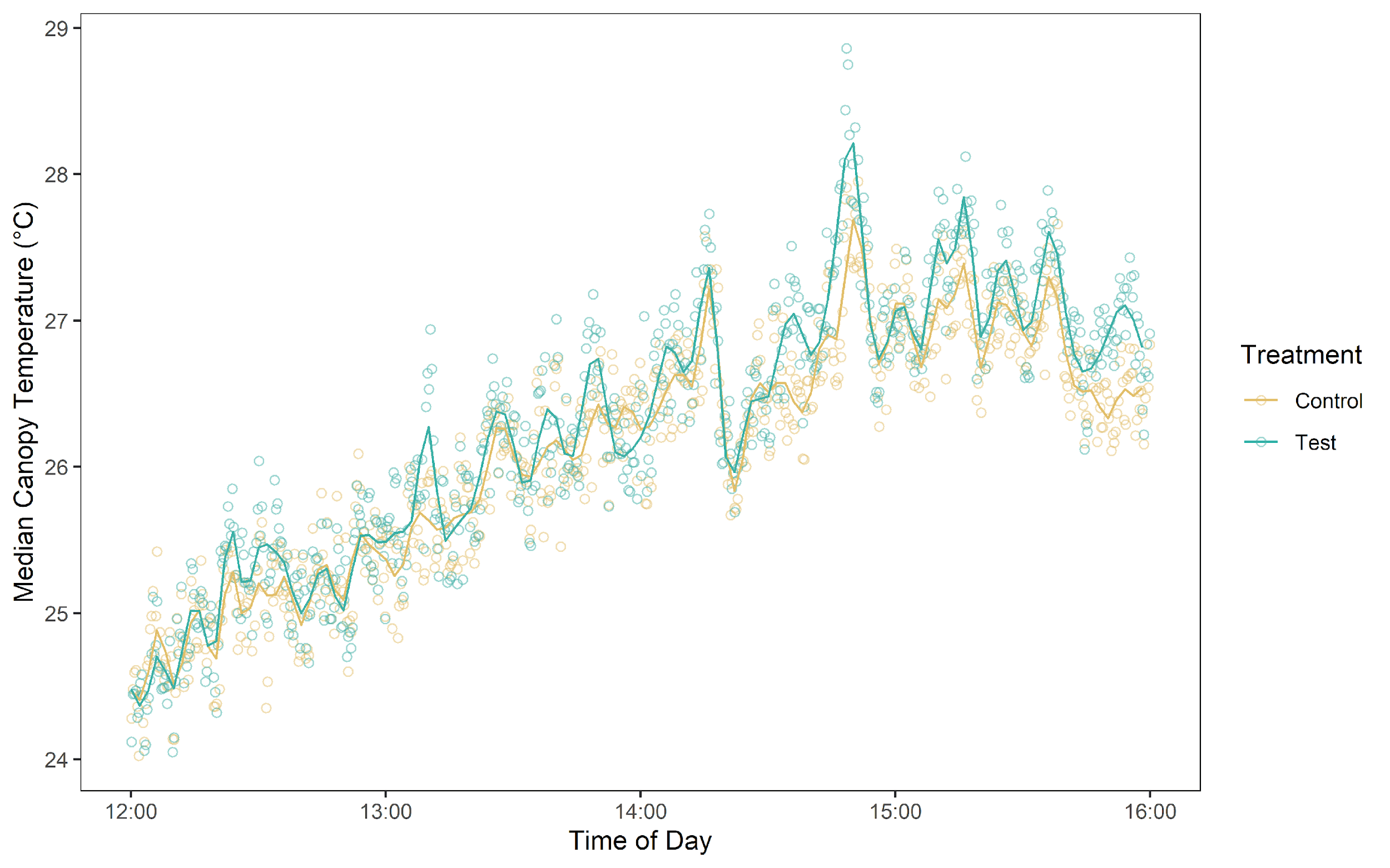

*Supplementary Figure S 3 Time courses of canopy temperature values during early afternoon on 8 June 2023 (233 DAS). Circles represent median canopy temperature values extracted from images taken every 20 s, the solid lines represent smoothed values from a spline fit. Data originates from two plots of the cultivar ‘Piznair’, growing side-by-side.*
